# Vitamin D receptor is necessary for metabolic health after sleeve gastrectomy

**DOI:** 10.1101/2025.10.24.684160

**Authors:** Andrei Moscalu, Weronika Stupalkowska, Lei Zhao, Yitong Li, Cullen F. Roberts, Yingjia Chen, Fei Ye, Rose Gold, Gavin Bewick, Francesco Rubino, Ali Tavakkoli, A. Sloan Devlin, Eric G. Sheu

**Author notes:** These authors contributed equally to this work (Andrei Moscalu, Weronika Stupalkowska, A. Sloan Devlin, Eric G. Sheu).

## Abstract

**Background:** The vitamin D receptor (VDR) regulates insulin sensitivity. Metabolic bariatric surgery (MBS) remains the most effective treatment for obesity and T2D. However, whether its metabolic effects are VDR-dependent is unknown. Here, we assessed VDR role in the metabolic response to sleeve gastrectomy (SG).

**Methods:** Whole body VDR knockout (KO) and wild type (WT) C57BL/6J mice with diet-induced obesity (DIO) were assigned to either SG or sham procedure. Postoperatively, animals underwent glucose and insulin tolerance tests. On sacrifice, serum, white adipose tissue (WAT), liver and colonic contents were collected for further biochemical, histological and bile acid analysis. Separately, human *VDR* gene expression was assessed in subcutaneous adipose tissue (SAT) biopsies collected from patients with/without MBS history.

**Results:** KO SG mice exhibited delayed glucose utilization after an oral challenge and progressive loss of insulin sensitivity, despite the same magnitude of surgery-induced weight change between the two genotypes. WAT in KO SG mice had lower mass, smaller adipocytes and increased inflammation. SG had a differential effect on colonic levels of the glucoregulatory cholic acid 7-sulfate (CA7S): increasing CA7S concentration in the WT mice but decreasing it in the KO mice. Finally, patients with MBS history had higher *VDR* expression in SAT as compared to those without MBS history.

**Conclusion:** VDR is necessary for metabolic health after SG in DIO mice due to its role in WAT function, insulin sensitivity and inflammatory response after surgery. Increased expression of VDR in patients post-MBS suggests that it may also contribute to metabolic responses in humans.

**Highlights:**

- VDR KO results in delayed glucose utilization following SG in DIO mice
- VDR KO results in progressive loss of improved insulin sensitivity after SG
- VDR KO results in white adipose tissue inflammation and remodeling after SG
- VDR expression increases in human subcutaneous adipose tissue after bariatric surgery

## 1 INTRODUCTION

Vitamin D receptor (VDR) mediates actions of the physiologically active 1,25-dihydroxyvitamin D3 (1). VDR is expressed throughout the human body, including in insulin-responsive tissues and organs such as adipose, muscle, liver and pancreas (2). Vitamin D-VDR signaling is implicated in a variety of biological functions that extend beyond the conventional calcium homeostasis and bone health. The extra-skeletal effects of vitamin D-VDR signaling include glucose and lipid metabolism (3), but the exact pathways are not well understood. Low levels of vitamin D are associated with obesity, insulin resistance (IR) and higher glucose levels (4, 5). However, studies examining the effects of vitamin D supplementation have yielded inconsistent results: while some authors report significantly improved insulin sensitivity and fasting glucose levels (6, 7), others did not record similar benefits (8–10). Supplementation of vitamin D resulted in increased VDR expression and decreased adiposity (11). In contrast, overexpression of human VDR in mouse adipose tissue (AT) led to obesity, IR and worsened glucose tolerance (12). Moreover, VDR can promote IR *in vitro* in mouse adipocytes (13). Overall, the inconsistent data on the role of VDR in metabolic health and the lack of any studies examining VDR signaling after metabolic bariatric surgery (MBS), represent a knowledge gap which this study aims to address.

It is important to note, that although the primary endogenous ligand for VDR is the active form of vitamin D (1,25-dihydroxyvitamin D3), certain bile acids (BAs) such as lithocholic acid (LCA) have also been reported to bind to VDR (14). In our prior work, we showed that decreased LCA in the gut post MBS resulted in increased LCA transport to the liver, where it activated VDR, resulting in increased expression of sulfotransferase enzyme (SULT2A1) and increased production of glucoregulatory BA cholic acid 7-sufate (CA7S) (15, 16). Together, these data suggest that signaling pathways involving VDR are complex and play an important role in regulation of metabolism. Further elucidation of the impact of VDR signaling on host metabolism could help pave the way for targeting VDR to treat metabolic diseases.

Sleeve gastrectomy (SG) is one of the most common MBS procedures performed worldwide (17). Recently, the effects of SG have been investigated in an animal model of diet-induced obesity (DIO) combined with vitamin D deficiency (VDD) (18). The authors concluded that VDD attenuated the benefits of surgery on IR and body weight (BW), with associated intestinal dysbiosis (18). Here, we took an alternative approach and characterized the post-surgery phenotypes in the whole body VDR knockout (KO) mice, hypothesizing that functional VDR is necessary for metabolic benefits after SG. We found that while VDR was dispensable for weight loss, it was required for optimal glucose utilization, maintaining insulin sensitivity, AT health and lipid homeostasis after surgery.

## 2 METHODS

### 2.1 Animals

All animal experiments were approved by the Institutional Animal Care and Use Committee at Brigham and Women’s Hospital (Boston, MA, USA). 3-week-old male C57BL6/J mice were purchased from Jackson Laboratory (Strain # 000664) and used as wild type (WT) control group. Three breeding pairs of whole body VDR KO mice (19) were generously donated by Dr Marie Demay (Massachusetts General Hospital, Boston, MA, USA) and used to establish a KO colony. The presence of VDR KO mutant sequence in the first-and second-generation mice was confirmed with RT-PCR (Transnetyx Inc). Data presented here are pooled from one cohort of WT mice and two cohorts of KO mice (two separate litters, operated on approximately one month apart).

Mice were housed in a climate-controlled environment with 12-hourly light-dark cycles. From 3 weeks of age, animals were fed high fat diet (HFD, 60% calories from fat) enriched in calcium (2%), phosphorus (1.25%) and lactose (20%) (D22071502, Research Diets; Table S1) until sacrifice at 27 weeks (15 weeks postoperatively), except for a brief perioperative period when they were administered gel diet (DietGel Recovery, Clear H2O) for 2 days before surgery and 5 days after.

### 2.2 Surgery

A previously established animal model of SG was used (20). At 12 weeks of age, following an overnight fast, mice underwent either SG or a sham procedure (SH). SG and SH groups within each genotype were matched by BW. All procedures were performed by the same surgeon (A.M.), in an alternating sequence, and on a heating pad under isoflurane anesthesia. After shaving, the anterior abdominal wall was disinfected, and an upper midline incision was made to access intraperitoneal cavity. The stomach was delivered into the wound and short gastric vessels were divided with electrocautery. For SG procedure a 45mm linear tri-stapler (Medtronic, New Haven, CT, USA) was used to remove approximately 80% of the stomach and create a small gastric tube. For SH procedure, only division of short gastric vessels was performed, followed by returning of the stomach into the abdominal cavity. Intraperitoneal 0.9% sodium chloride was administered to maintain adequate hydration. Abdominal wall fascia and skin were closed in layers with interrupted 6/0 Vicryl sutures (Ethicon Inc., Raritan, NJ, USA). Subcutaneous extended-release buprenorphine was used for postoperative analgesia.

### 2.3 Postoperative monitoring

Activity, pain level, incision and body condition were monitored twice daily for the first 5 days after surgery. BW was measured daily from postoperative day 1 until day 7, weekly until day 63, and every 2-4 weeks thereafter. Food intake was measured for each mouse individually every few days for approximately 9 weeks postoperatively.

### 2.4 Glucose tolerance and insulin tolerance tests

Oral glucose tolerance tests (OGTTs) were conducted at postoperative weeks 2 and 8. Insulin tolerance tests (ITTs) were conducted at postoperative weeks 5 and 9. Mice were fasted for 4 hours. After baseline fasting glucose measurement, animals were administered either 2g/kg glucose (Teknova) by oral gavage (OGTT) or 0.6U/kg insulin (Eli Lilly) via intraperitoneal injection (ITT). Glucose measurements were taken at 15,30,60 and 120 minutes (OGTT) or 15,30,45,60,90 and 120 minutes (ITT) with OneTouch Ultra 2 glucometer (LifeScan).

### 2.5 Serum calcium and lipid profile

Serum samples were collected at sacrifice after an overnight fast. Serum calcium concentration was determined using Total Calcium LiquiColor kit (EKF Diagnostics). Serum cholesterol, high-density lipoprotein (HDL) and triglycerides were measured with CardioChek PA Analyzer (PTS Diagnostics).

### 2.6 Histology

Liver, epididymal white adipose tissue (eWAT) and inguinal white adipose tissue (iWAT) specimens were stained with hematoxylin and eosin (H&E) and liver slides were additionally stained with Oil Red O (ORO) and Sirius Red (SRS) (Histology Core, Beth Israel Deaconess Medical Center, Boston, MA, USA). Photomicrographs were taken at 10x magnification (Olympus BX50 microscope, QiCam digital camera, Image-Pro Premier software). For eWAT and iWAT, 1-3 photomicrographs were randomly taken from each H&E slide for quantitative analysis. Adipocyte counting and surface area estimation was performed using Adiposoft plugin (21) for ImageJ2 (Fiji). Furthermore, eWAT and iWAT slides were reviewed by a clinical pathologist (L.Z.) who was blinded to the intervention groups. The pathologist scored inflammation and fat necrosis using a customized scale (Table S2) and assessed for presence of fibrosis. Additionally, the same pathologist reviewed liver slides evaluating steatosis, inflammation and fibrosis (22). Other investigators were not blinded during the experiment conduct or data analysis.

### 2.7 BAs analysis

BA analyses were performed using a previously reported method (23). Briefly, colon contents (30–100 mg) were weighed, and 500 µL of HPLC-grade methanol containing 100 µM glycochenodeoxycholic acid (internal standard) was added to each homogenization vial. Samples were homogenized using an OMNI Bead Mill Homogenizer at 6.3 m/s for 16 minutes to ensure thorough disruption. Following homogenization, the vials were centrifuged at 20,000 × g for 15 minutes. A 100 µL aliquot of the supernatant was collected and mixed with an equal volume of 50% methanol before analysis. BAs quantification was performed using an Ultra-high Performance Liquid Chromatography-Mass Spectrometry (UPLC-MS) system following established protocols (23). Calibration curves for each BA were determined according to previously published procedures, and detection limits were the same as previously described (16).

### 2.8 Animal exclusions

Mice that did not survive until the study endpoint or were found to have a leak on necropsy were excluded from all analyses (WT SH= 0, WT SG = 3, KO SH = 4, KO SG = 12). Additionally, one WT SG mouse, one KO SG mouse and two KO SH mice were selectively excluded from histological analyses due to either incidental anatomical anomalies unrelated to surgery or planned sacrifice before study endpoint. Another two KO SH mice in eWAT analysis and one KO SH mouse in iWAT analysis were excluded due to lack of corresponding HE slides. One WT SG mouse was selectively excluded from 2-week OGTT data analysis due to failure of oral gavage.

### 2.9 Human samples and data

Institutional Review Board approval was granted to collect tissue samples and gather clinical data from patients undergoing elective abdominal surgery in the Department of Gastrointestinal Surgery, Brigham and Women’s Hospital (Boston, MA, USA). All patients signed written consent forms.

SAT biopsies were obtained from laparoscopic port sites, transported on ice and stored at −80°C. From available samples we identified 11 patients with history of MBS and BMI below 40 kg/m^2^ at the time of biopsy and 9 control patients without MBS history. Patients were matched by age, sex, race, ethnicity, diabetes status, BMI and vitamin D level.

### 2.10 RNA extraction from human SAT and qPCR

Approximately 150 mg of SAT was used to extract total RNA (RNeasy Lipid Tissue Mini Kit, Qiagen). Following reverse transcription (Reverse Transcription Kit, Applied Biosystems) RT-qPCR was performed (SYBR Green; Applied Biosystems) using the Quant Studio 6 (ThermoFisher). The 2^-ΔΔCt^ method was used to calculate the relative change in gene expression. *HSPCB* gene was used for normalization (24). Primer sequences are listed in Table S3.

### 2.11 Statistical analysis

Statistical analyses were performed using GraphPad Prism 10.3.0 and Microsoft Excel. All data are presented as mean ± SEM unless stated otherwise. Statistical significance threshold was set at p<0.05. Continuous data were compared either with two-tailed Student’s t-test (two groups), one-way ANOVA (three groups) or two-way ANOVA with Sidak’s multiple comparisons test (four groups). For repeated measures four group comparison a two-way repeated measures ANOVA with Tukey’s multiple comparisons test was used. For *post hoc* multiple comparisons the following pairwise comparisons were pre-selected: WT SH vs WT SG, KO SH vs KO SG, WT SH vs KO SH and WT SG vs KO SG. For categorical data comparisons, Fisher’s exact test was used.

## 3 RESULTS

### 3.1 VDR is dispensable for SG-induced weight loss

We utilized our previously established model of SG (20) in DIO mice to characterize the phenotype of SG in the absence of VDR (Fig 1A). Whole body VDR KO was previously described as resistant to DIO (25, 26) but in our hands, KO mice were not protected from obesity when fed 60% HFD. In fact, preoperatively, the first cohort of KO mice was significantly heavier than the WT group (Fig S1A). Postoperatively, SG induced weight loss in both WT and KO animals (Fig 1B with Table S2; and Fig S1B-E with Table S3). Both KO SH and KO SG gained less weight postoperatively as compared to their WT counterparts (Fig 1B, Table S2). However, when the mean weight change in SG was expressed as a ratio of mean weight change in SH, the degree of weight loss induced by SG did not differ between genotypes except at postoperative week 1 and on necropsy day which both included fasting periods (Fig 1C and Fig S1F). While KO mice exhibited marginally lower food intake compared to WT mice, SG did not affect food intake in either genotype apart from postoperative week 1 (Fig 1D, Table S4). Overall, these data suggest that VDR is dispensable for SG-induced weight loss in DIO mice.

**Figure 1.**
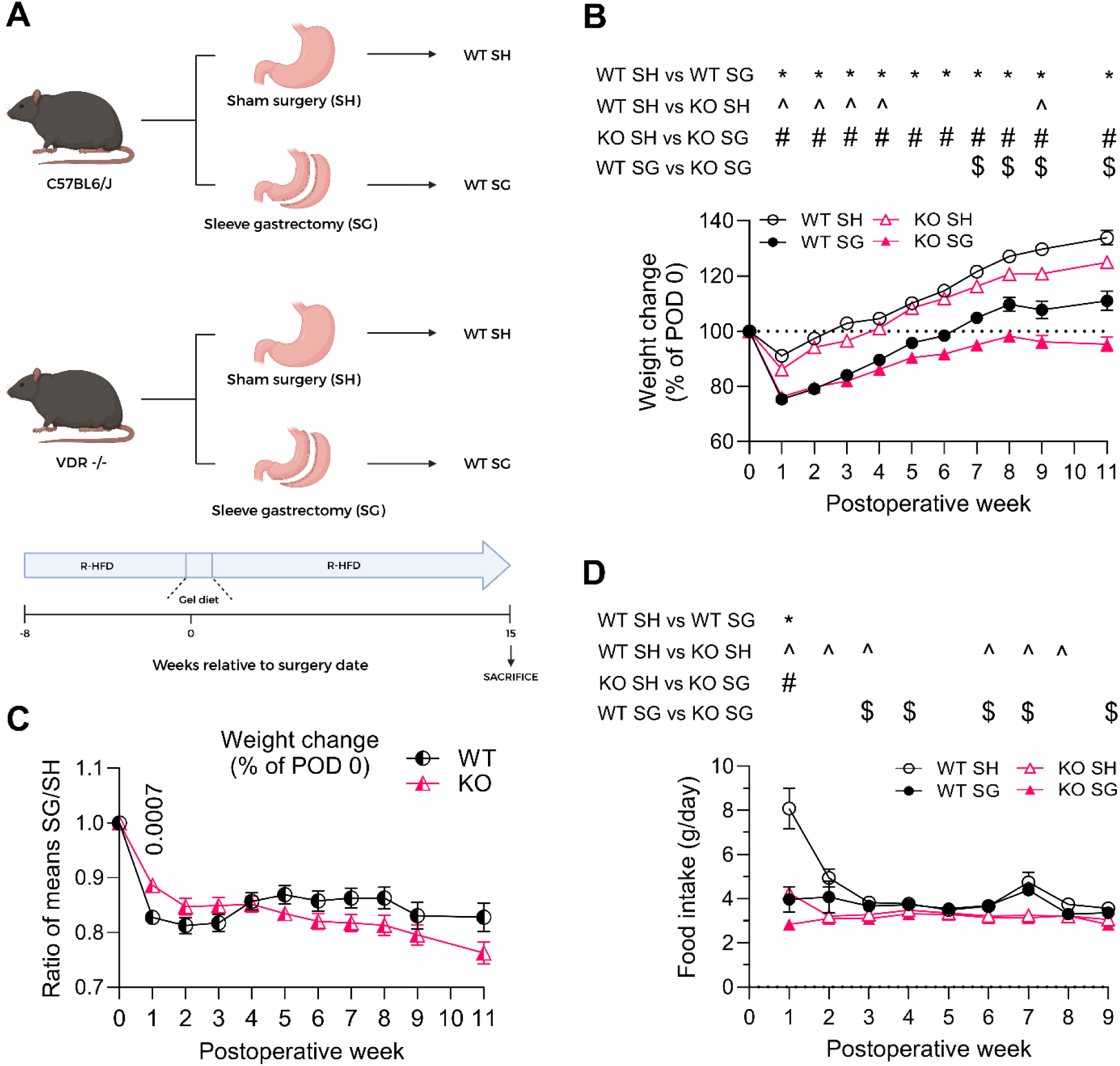
VDR is dispensable for SG-induced weight loss. **(A)** Experimental design. Whole body VDR knockout (KO) and wild type (WT) C57BL/6J male mice were fed calcium-phosphate-lactose supplemented high-fat (60%) diet (R-HFD). At the age of 12 weeks, mice were randomly assigned to either sleeve gastrectomy (SG) or sham procedure (SH) and data was recorded for 15 weeks postoperatively. **(B)** Change in body weight over time as % of body weight on postoperative day 0 (POD 0). **(C)** SG/SH ratio of mean weight change over time per genotype. **(D)** Average daily food intake over time. Animal numbers: WT SH (n=8), WT SG (n=9), KO SH (n=21), KO SG (n=21). Data in panels B and D are shown as means ±SEM, panel C shows ratio of mean SG/SH weight change ± SE. Data in panels B and D were analyzed with two-way repeated measures ANOVA with Tukey’s multiple comparisons test. Corresponding p values are given in Supplementary Tables 2 and 4. In panel C, a Student’s t test was performed for each postoperative week. Data not marked are not significant.

### 3.2 Lack of VDR results in delayed glucose utilization after SG

We performed the first OGTT two weeks postoperatively and assessed total areas under curve (tAUC) (Fig 2A-C). The tAUC revealed that SG improved glucose tolerance regardless of VDR status (Fig 2A; reduction in 2-week OGTT tAUC: WT −36.9%, p<0.0001; KO −36.1%, p<0.0001). The degree of improvement in glucose tolerance, when expressed as a ratio of mean tAUC in SG over mean tAUC in SH, did not differ between genotypes (Fig 2C, Fig S2A). However, glucose utilization after an oral challenge was altered in KO SG mice. Specifically, we observed delayed decline in blood glucose levels in KO SG compared to WT SG mice (Fig 2A, B). In the latter group, glucose levels peaked at 15 minutes post oral challenge and then decreased almost back to baseline by the 30-minute timepoint (Fig 2A, B). In contrast, in KO SG mice, glucose levels remained persistently elevated for the first 30 minutes and returned to baseline by the 60-minute timepoint (Fig 2A, B; Fig S2A).

**Figure 2.**
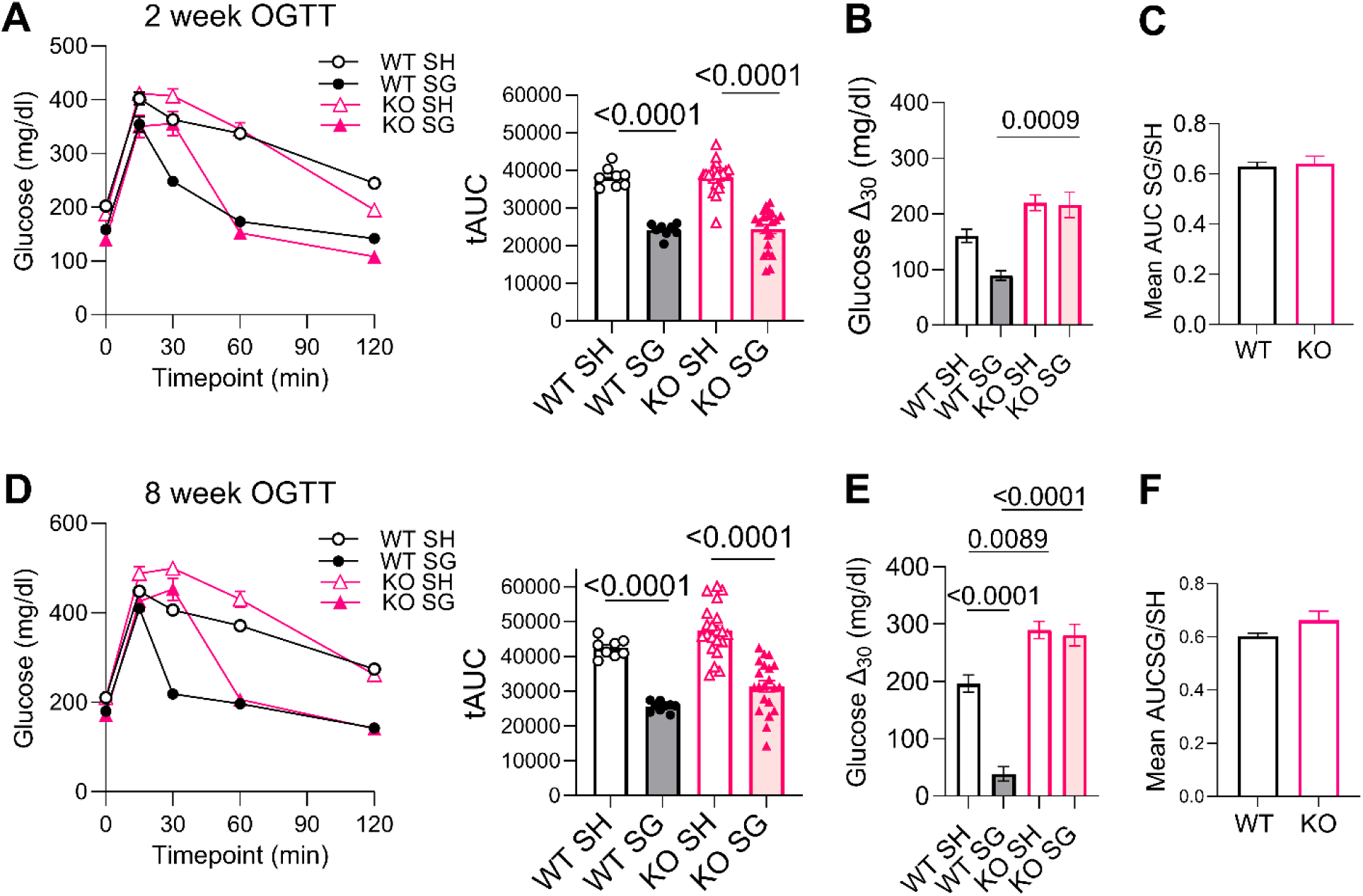
Lack of VDR results in altered glucose handling after SG. **(A)** 2-week oral glucose tolerance test (OGTT) with total Area Under Curve (tAUC) per experimental group. **(B)** Change in glucose between baseline and 30 minutes during the 2-week OGTT (Δ_30_). **(C)** SG/SH ratio of mean tAUC for 2-week OGTT per genotype. **(D)** 8-week oral glucose tolerance test with tAUC per experimental group. **(E)** Change in glucose between baseline and 30 minutes during the 8-week OGTT (Δ_30_) **(F)** SG/SH ratio of mean tAUC for 8-week OGTT per genotype. Animal numbers: WT SH (n=8), WT SG (n=9), KO SH (n=21), KO SG (n=21). In panel A one WT SG mouse was excluded from the analysis due to failure of oral gavage. Data in A, B, D and E are shown as mean ± SEM. C and F show ratios of SG/SH tAUC means ± SEM. Data were analyzed either with two-way ANOVA with Sidak’s multiple comparisons test (A, B, D, E) or two-tailed Student’s t test (C, F). Data not marked are not significant.

The SG-induced improvement in glucose tolerance was again observed in both genotypes at eight weeks postoperatively (Fig 2D-F; reduction in 8-week OGTT tAUC: WT −39.7%, p<0.0001; KO −33.7%, p<0.0001). Similar to 2-week OGTT, there was no effect of SG on reducing glucose levels within the first 30 minutes post-challenge in the KO mice (Fig 2E, Fig S2B). Overall, these data suggest that VDR signaling contributes to improved early-phase glucose utilization after surgery.

### 3.3 Lack of VDR results in progressive loss of insulin sensitivity after SG

Five weeks after surgery, we observed an improvement in insulin sensitivity in both KO SG and WT SG mice (Fig 3A,B; reduction in 5-week ITT tAUC: WT −29.3%, p=0.0339; KO −25.2%, p=0.0025). Notably, however, nine weeks post-surgery, KO SG mice no longer demonstrated improved insulin sensitivity compared to KO SH mice and in contrast to continued sensitivity in WT SG mice (Fig 3C, D; reduction in 9-week ITT tAUC: WT −33.2%, p=0.0276; KO −9.5%, not significant). This effect was consistent across both KO cohorts (Fig S3) and occurred despite the reduced BW in SG mice within each genotype. Overall, these data suggest that VDR is necessary to preserve improved insulin sensitivity after SG.

**Figure 3.**
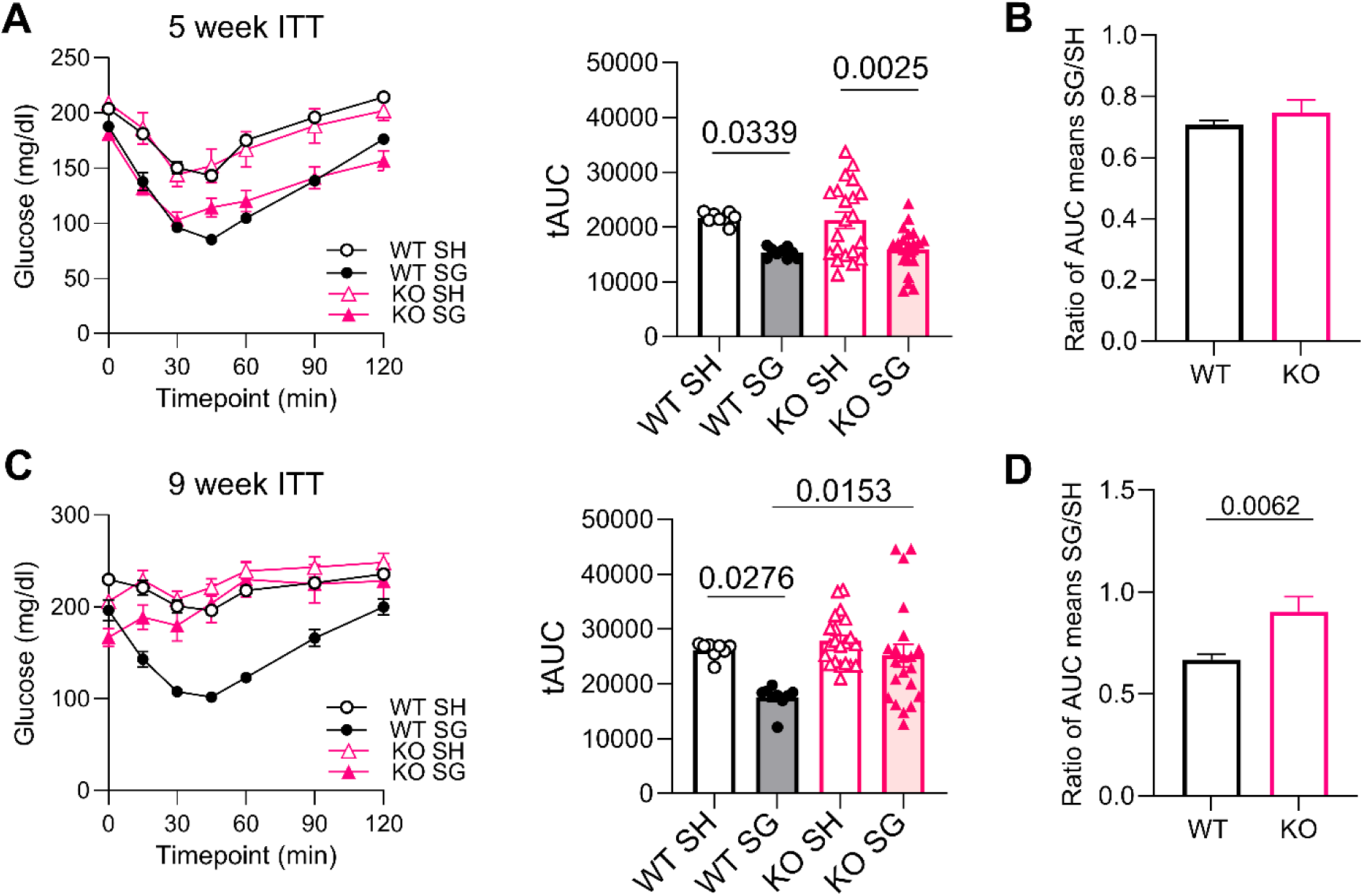
Lack of VDR results in progressive loss of insulin sensitivity after SG. **(A)** 5-week insulin tolerance test (ITT) with total Area under Curve (tAUC) per experimental group. **(B)** SG/SH ratio of mean tAUC for 5-week ITT per genotype. **(C)** 9-week ITT with tAUC per experimental group. **(D)** SG/SH ratio of mean tAUC for 9-week ITT per genotype. Animal numbers: WT SH (n=8), WT SG (n=9), KO SH (n=21), KO SG (n=21). Data in A and C are shown as mean ± SEM. B and D show ratios of SG/SH tAUC mean ± SEM. Data in A and C were analyzed with two-way ANOVA with Sidak’s multiple comparisons test. Data in B and D were analyzed with two-tailed Student’s t test. Data not marked are not significant.

### 3.4 VDR is necessary to maintain healthy AT after SG

Upon animal sacrifice, it became evident that KO SG mice have macroscopically smaller WAT depots. The mass of iWAT, normalized to BW, was lower in SG mice regardless of genotype and numerically lowest in KO SG mice (Fig 4A and Fig S4A). The normalized mass of eWAT was significantly reduced only in KO animals which was mainly driven by the second cohort of KO mice (Fig 4A and Fig S4B).

**Figure 4.**
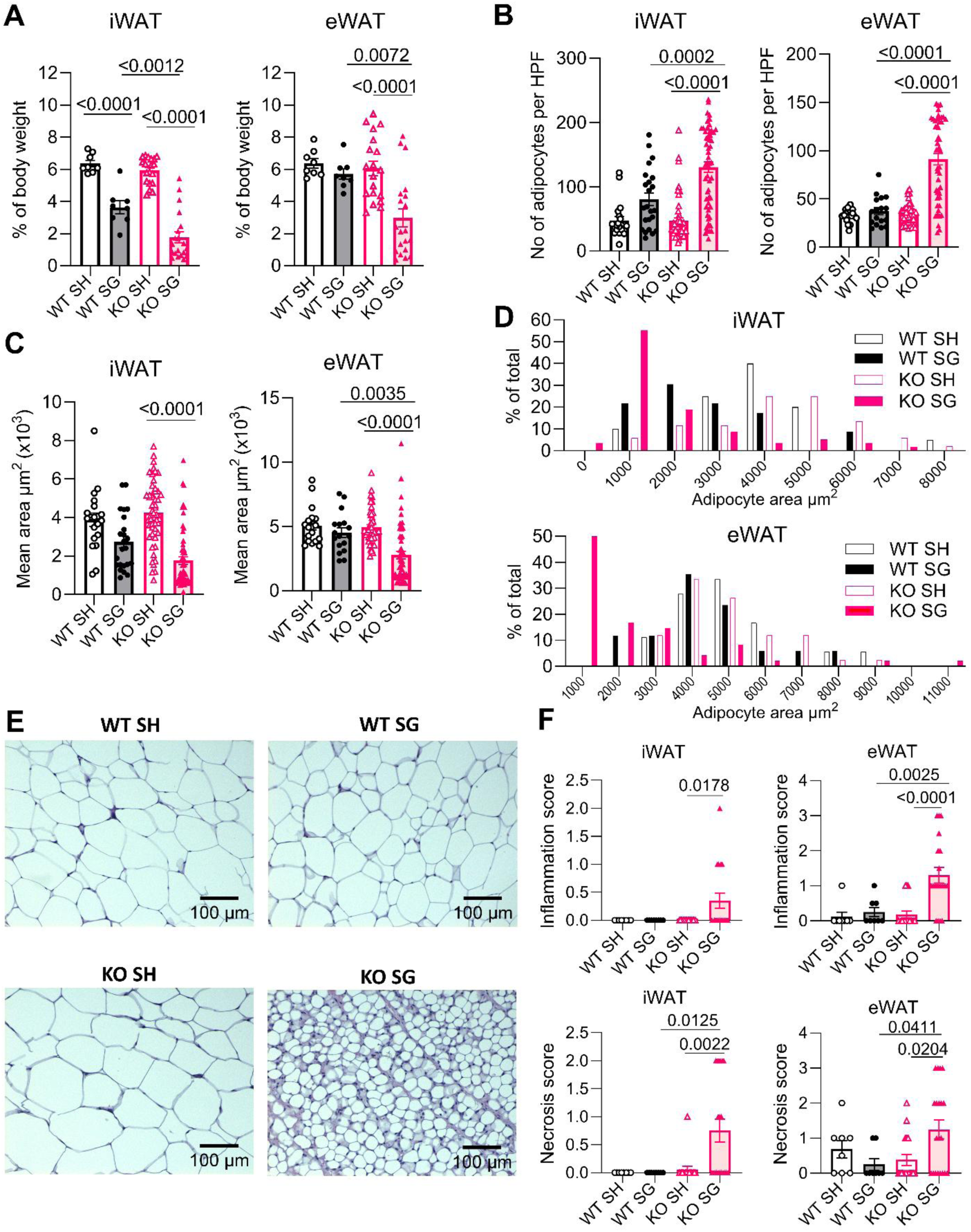
VDR is necessary to maintain healthy adipose tissue after SG. **(A)** Mass of inguinal (iWAT) and epididymal (eWAT) white adipose tissue depots as percentage of body weight on necropsy day. **(B)** Number of adipocytes per x10 high power field (HPF) in iWAT and eWAT. **(C)** Mean adipocyte area in iWAT and eWAT. **(D)** Percentage distribution of adipocyte area among experimental groups. **(E)** Representative photomicrographs at x10 magnification of iWAT stained with hematoxylin & eosin (H&E). **(F)** Inflammation and adipocyte necrosis scores (0–3) in iWAT and eWAT. Animal numbers: WT SH (n=8), WT SG (n=8), KO SH (n=19), KO SG (n=20). In panel A each symbol represents an individual mouse. In panels B-C each symbol represents an individual photomicrograph (for each mouse, one H&E slide was produced per adipose tissue type and 1-3 representative photomicrographs were taken of each slide for automated analysis). Panel D shows percentage of photomicrographs with mean adipocyte area in predetermined size bins. In panel F each symbol represents an individual H&E slide (one per mouse). Data shown as mean ± SEM. Data in panels A-C and F were analyzed with two-way ANOVA with Sidak’s multiple comparisons test. Data not marked are not significant.

Further histological analysis revealed that morphology of both iWAT and eWAT in KO SG mice was strikingly different than in the other experimental groups. AT depots from KO SG mice had a significantly higher number of adipocytes per high power field (Fig 4B and Figure S5A,B). Moreover, the mean adipocyte area was significantly lower in KO SG mice (Fig 4C and Fig S5C,D). 59% of iWAT and 50% of eWAT adipocytes fell into the smallest size bin in the KO SG group as compared to WT mice where the distribution of adipocyte size followed a more bell-shaped curve regardless of surgery type (Fig 4D). Representative photomicrographs demonstrate these striking morphological differences (Fig 4E).

Alterations in adipocyte size and cellularity were not the only differences which set the KO SG group apart. We noticed significant inflammatory infiltrate in both iWAT and eWAT of KO SG mice (Fig 4F). These changes were accompanied by increased fat necrosis (Fig 4F). Both inflammation and fat necrosis appeared to be slightly more prominent in the second KO cohort (Fig S6). Finally, two of the KO SG mice had fibrosis in their WAT depots, as compared to none in the other experimental groups. The KO SG photomicrograph in Fig 4E represents a fibrotic response in iWAT. Overall, these data suggest that both iWAT and eWAT undergo substantial remodeling in KO SG mice, including reduced total mass, reduced size of adipocytes, and increased inflammation and fat necrosis.

### 3.5 VDR contributes to lipid and calcium homeostasis after SG

The morphological changes in WAT of KO SG mice may suggest a deficiency in lipid storage function and consequent diversion of lipids to other sites, such as liver and circulation. We therefore assessed hepatic lipid storage and serum lipid profile. The absolute liver mass was reduced by SG regardless of genotype (Fig 5A and Fig S7A). However, the normalized liver mass was only reduced in WT SG mice (Fig 5A and Fig S7B). Given that the normalized mass of WAT is lowest in KO SG and normalized mass of liver is numerically lowest in WT SG, these data demonstrate a differential effect of SG on reduction in mass of different organs across genotypes. WT animals lose relatively more liver mass, whereas KO animals lose relatively more WAT mass.

**Figure 5.**
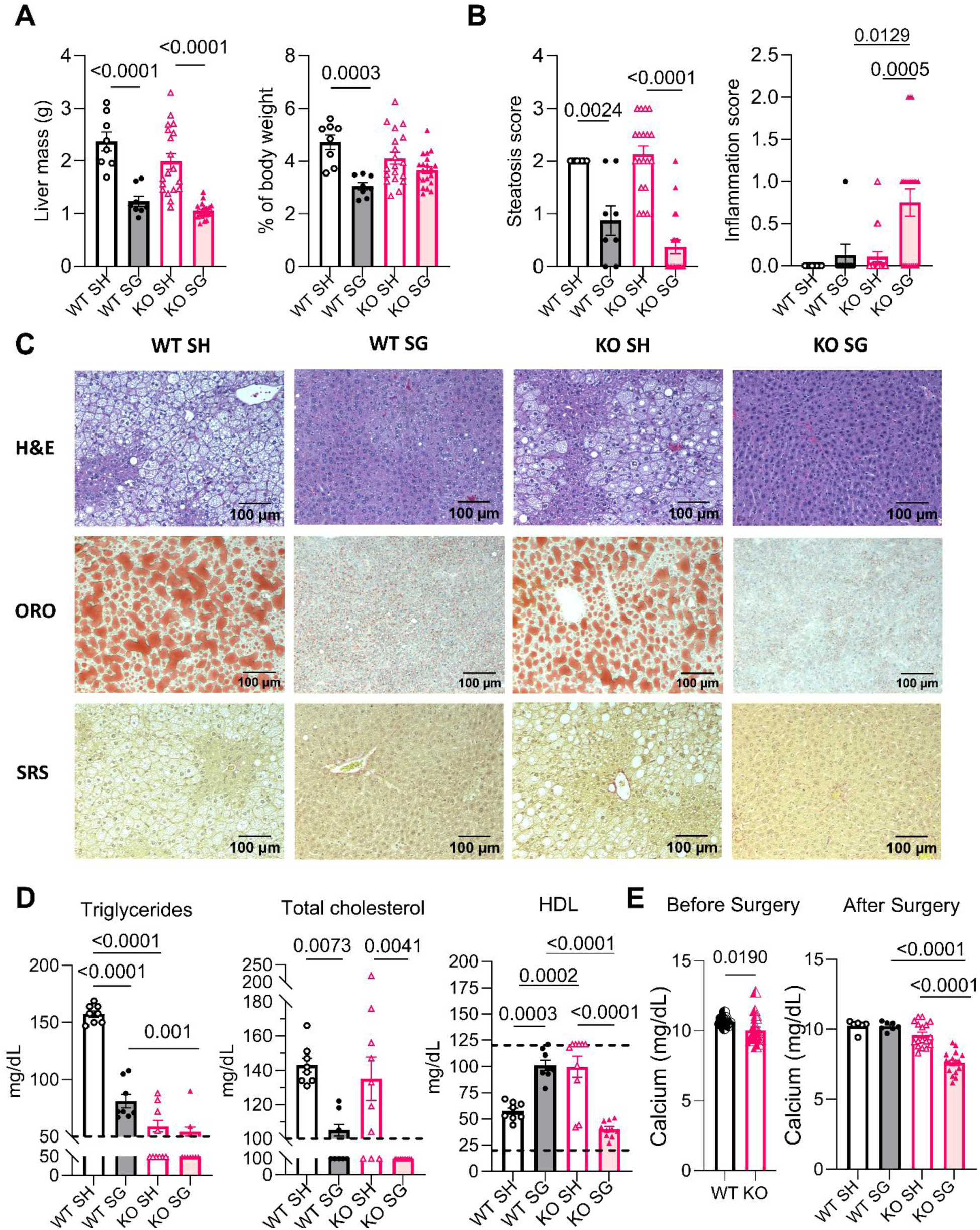
VDR contributes to lipid and calcium homeostasis after SG. **(A)** Liver mass (left) and liver mass as percentage of body weight (right) on necropsy day. **(B)** Hepatic steatosis and inflammation scores (0-none, 1-mild, 2-moderate, 3-severe). **(C)** Representative photomicrographs of liver sections in each experimental group stained with hematoxylin & eosin (H&E; top panel), Oil Red O (ORO; middle panel), and Sirius Red (SRS, bottom panel). Taken at x10 magnification. **(D)** Postoperative serum lipid profile. Dashed lines represent the lower and upper (when within the graph range) detection limits of the assay (triglycerides: 50-500 mg/dL; cholesterol: 100-400 mg/dL; HDL: 20-120 mg/dL). Datapoints falling outside the detection range are marked at the lower or upper limit as applicable. **(E)** Preoperative (left) and postoperative (right) levels of serum calcium in WT and KO mice measured at 7 weeks of age (preoperative) and 27 weeks of age (postoperative). Animal numbers (A-B): WT SH (n=8), WT SG (n=8), KO SH (n=19), KO SG (n=20). Animal numbers in D: WT SH (n=8), WT SG (n=8), KO SH (n=10), KO SG (n=10). Animal numbers in E: WT (n=12), KO (n=18), WT SH (n=6), WT SG (n=7), KO SH (n=17), KO SG (n=17). Each symbol represents an individual mouse. Data shown as mean ± SEM. Data were analyzed with two-way ANOVA with Sidak’s multiple comparisons test. Data not marked are not significant.

On histological assessment, hepatic steatosis was present in both WT SH and KO SH mice, and SG reduced steatosis in both genotypes (Fig 5B,C and Fig S7C). In addition, similarly to postoperative changes seen in WATs, there was an increased inflammation in the livers of KO SG mice (Fig 5B and Fig S7D). Hepatic fibrosis was noted in four KO SG mice, and it was absent in the other experimental groups (p=0.1230; Table S5). Overall, these data suggest that VDR is dispensable for SG-induced reduction in steatosis, but similar to its role in WAT, it regulates hepatic immune response after surgery.

KO SH mice had significantly lower serum triglyceride levels than WT SH (Fig 5D), consistent with prior reports in surgery-naïve WT and KO mice (25). In fact, the level of triglycerides in a high proportion of both KO SH and KO SG mice was below the detection limit, therefore it was impossible to assess the effects of surgery on the triglyceride level in KO mice. Furthermore, in all KO SG and many WT SG mice, the levels of cholesterol were also below the detection limit (Fig 5D). Nevertheless, it is possible to conclude that VDR is dispensable for surgery-induced reduction in serum cholesterol. Finally, with regards to HDL, a significant proportion of KO SH mice had HDL level above the detection limit and much higher than WT SH mice (Fig 5D). There was a striking difference in the effect of surgery on HDL levels across genotypes: in WT mice, SG increased HDL but in KO mice, surgery resulted in significantly lower HDL levels (Fig 5D). Of note, SG decreased levels of total serum calcium in KO animals, despite the calcium-supplemented diet (Fig 5E), which was previously shown to protect against hypocalcemia in surgery-naïve VDR KO mice (27). Overall, these data show that in addition to markedly reduced mass and altered morphology of WAT, SG in KO mice leads to reduction in hepatic lipid content, low levels of circulating lipids, and induction of hypocalcemia, suggesting that VDR plays an important role in regulating lipid and calcium homeostasis after SG.

### 3.6 Lack of VDR results in decreased colonic cholic acid 7-sulfate and increased β-muricholic acid after SG

The total BAs level in colonic contents of KO mice was numerically higher as compared to WT animals (Fig 6A). This result is consistent with existing literature showing that naïve whole body VDR KO mice possess higher enterohepatic BA concentrations (28). SG did not alter total BA concentration in either genotype (Fig 6A). In WT animals, consistent with our prior work (15), we observed numerically higher levels of the glucoregulatory BA cholic acid 7-sulfate (CA7S) and lower levels of LCA after SG (Fig 6B and Fig S8). The level of CA7S was higher in KO SH compared to WT SH animals. However, in KO mice, CA7S was the only individual BA that was significantly lower in KO SG as compared to KO SH (Fig 6B), which is consistent with the worsened glucose regulation observed in these animals. Finally, the only individual BA that was significantly higher in KO SG mice as compared to all the other experimental groups was beta-muricholic acid (βMCA) (Fig 6C). Overall, these data show that in the absence of VDR, SG decreases colonic CA7S levels, which may contribute to worsened glucose regulation observed in these animals.

**Figure 6.**
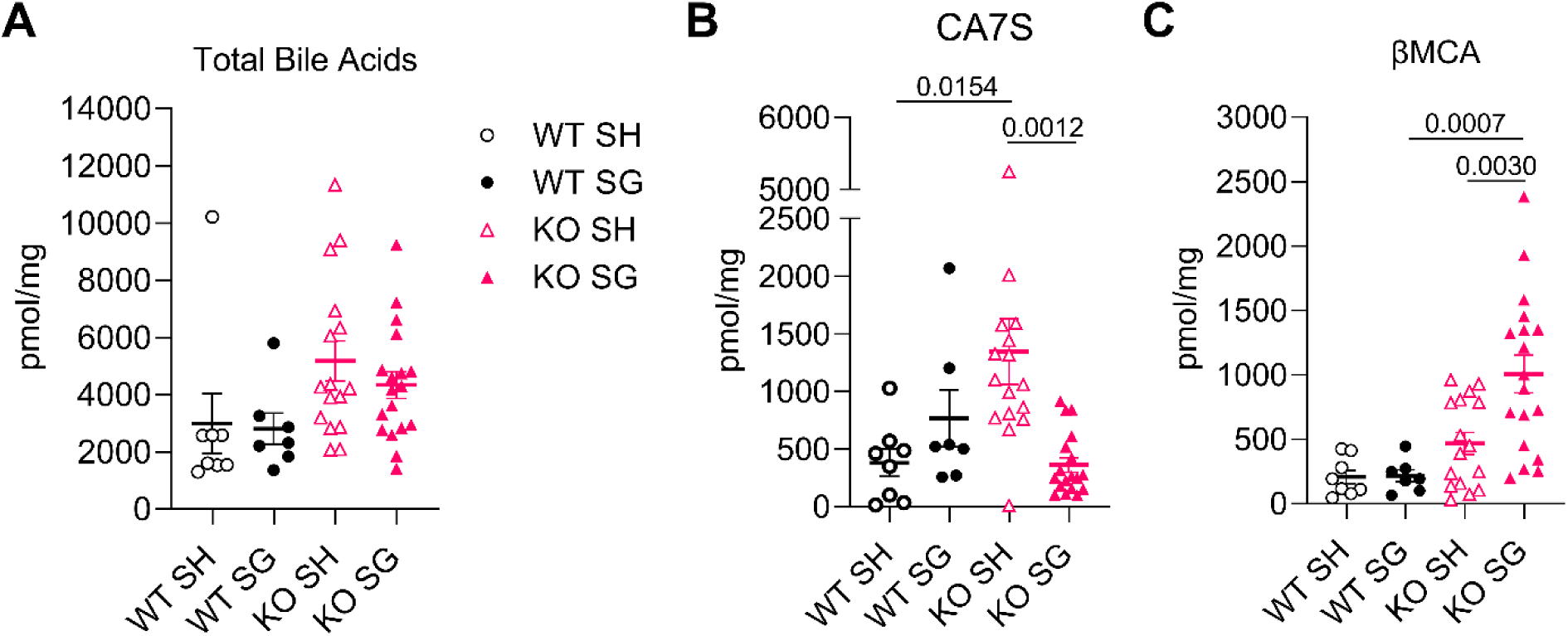
Lack of VDR results in decreased cholic acid 7-sulfate and increased β-muricholic acid after SG. **(A)** Total bile acids in colonic contents representing sum of all bile acids with measurable concentrations above limits of detection. **(B)** Colonic concentrations of bile acid CA7S per experimental group. **(C)** Colonic concentrations of bile acid β-MCA per experimental group. Animal numbers: WT SH (n=8), WT SG (n=7), KO SH (n=16), KO SG (n=18). Each symbol represents an individual mouse. Data presented as mean ± SEM. Data was analyzed with two-way ANOVA with Sidak’s multiple comparisons test. Data not marked are not significant. Total bile acids in panel A*: αMCA, alpha-muricholic acid; βMCA, beta-muricholic acid; CA, cholic acid; CA7S, cholic acid 7-sulfate; DCA, deoxycholic acid; isoDCA, isodeoxycholic acid; LCA, lithocholic acid; Tα/βMCA, tauro-alpha and tauro-beta-muricholic acid*; *TCA, taurocholic acid; TCDCA, taurochenodeoxycholic acid; TDCA, taurodeoxycholic acid; ωMCA, omega-muricholic acid; 3-oxo DCA / 3-oxo CDCA; 3-oxo deoxycholic acid and 3-oxo chenodeoxycholic acid; 3-oxo LCA, 3-oxo lithocholic acid; 7-oxo CA, 7-oxo cholic acid*.

### 3.7 MBS is associated with increased VDR expression in human SAT

To explore the role of VDR in response to MBS in humans, we obtained SAT biopsies from 11 patients with MBS history and 9 patients without MBS history who were matched by age, sex, race, ethnicity, diabetes status, BMI at the time of biopsy and serum vitamin D level within 1 year before the biopsy (Fig 7A-B). Patients with MBS history lost on average 20% of their total weight after a median of 5 years (Figure 7B). In MBS patients, we observed increased expression of *VDR* in SAT compared to individuals without MBS history (Fig 7C). These data suggest that MBS may induce *VDR* expression in human SAT. These results motivate future studies investigating whether increased VDR expression in post-MBS patients contributes to the long-term metabolic benefits of this procedure.

**Figure 7.**
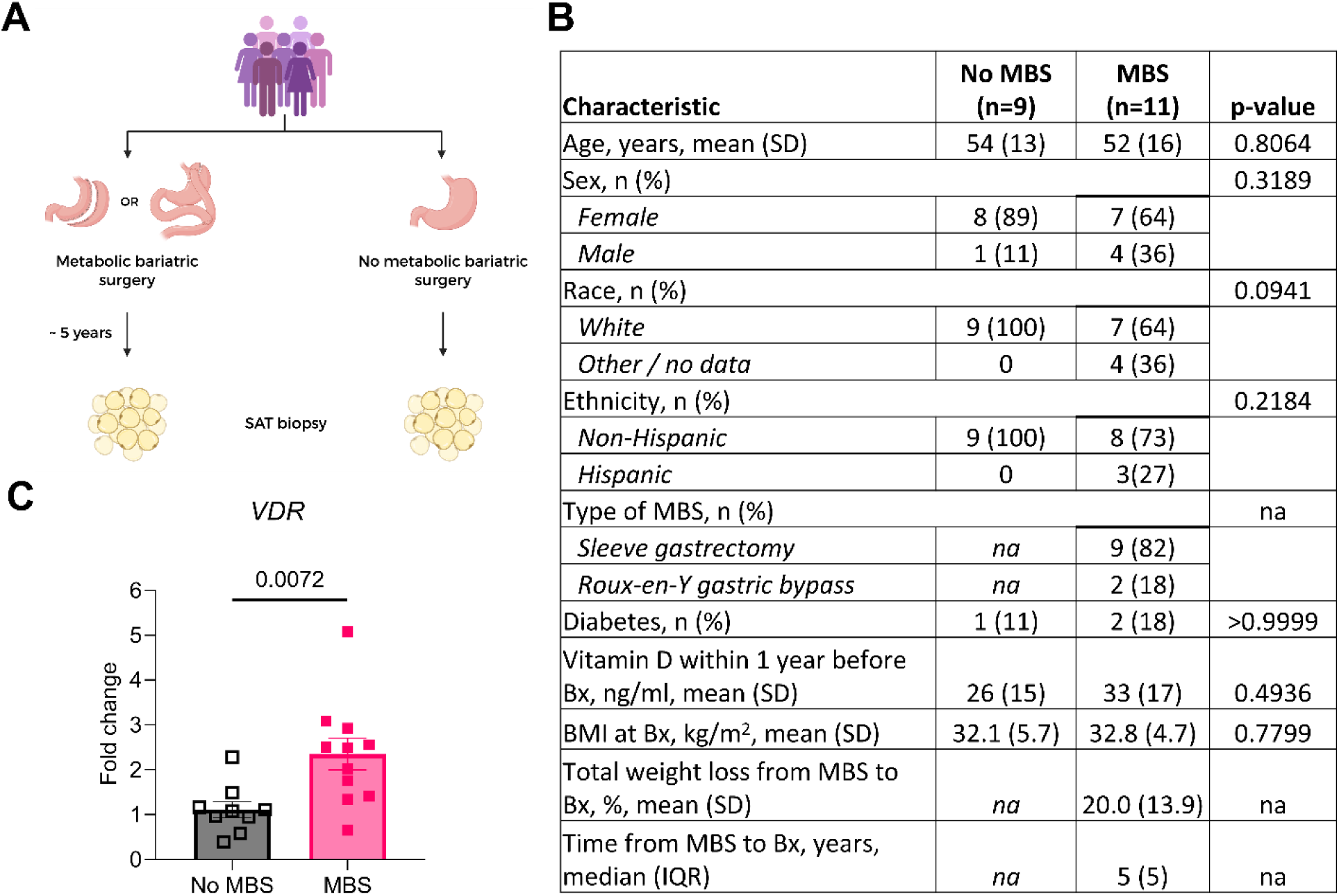
Metabolic bariatric surgery (MBS) is associated with increased VDR expression in human subcutaneous adipose tissue (SAT). **(A)** Experimental design. SAT biopsies were obtained from individuals either with (n=11) or without (n=9) history of MBS. **(B)** Baseline characteristics of patients whose SAT biopsies were used to assess *VDR* expression. **(C)** Relative expression levels of *VDR* gene in SAT biopsies from patients with (MBS) or without (no MBS) history of MBS. Between-group differences in B were tested either with Student’s t-test with Welch correction (continuous data) or Fisher’s exact test (categorical data). Data in C are presented as mean ± SEM and analyzed with Student’s t-test.

## 4 DISCUSSION

Here we show that VDR is necessary for metabolic health after SG in mice through its involvement in regulation of glucose utilization, insulin sensitivity, lipid profile, serum calcium levels, WAT biology and inflammatory response after surgery. SG in VDR KO mice leads to a unique phenotype of deficient WAT, hepatic and adipose inflammation, hypolipidemia, hypocalcemia, altered glucose utilization and progressive loss of insulin sensitivity despite surgically induced weight loss. Moreover, SG in KO mice has the opposite effect on the glucoregulatory CA7S as compared to WT mice – surgery leads to decreased colonic levels of CA7S in KO SG mice, which is consistent with the worsened glucose homeostasis observed in these animals.

In humans, VDD and certain VDR polymorphisms have been shown to be associated with IR (2, 8, 29, 30). Mechanistically, the vitamin D-VDR axis is involved in regulation of insulin signaling through induction of insulin receptor gene expression. VDR activation leads to upregulation of insulin receptors on the cell surface and increased insulin-dependent glucose uptake (29, 31, 32). Therefore, lack of VDR can lead to decreased insulin receptor expression and decreased glucose uptake in insulin-responsive tissues. However, in our model, VDR KO on its own was not sufficient to affect IR in DIO mice, as evidenced by lack of significant differences in ITT between the SH groups at either timepoint. Another consequence of VDR KO that could explain progressive loss of insulin sensitivity is inflammation (2, 7, 33). VDR signaling decreases expression of pro-inflammatory markers and leads to increased glucose uptake in adipocytes and improved hepatic insulin sensitivity (7, 34). Conversely, chronic, low-grade inflammation, very similarly to VDD, is causally linked to IR by decreasing insulin signaling via inhibition of insulin receptor and insulin receptor substrate (35). Taken together, we hypothesize that over time, lack of VDR leads to increasing inflammatory infiltrate in WAT of KO SG mice and consequently these two factors contribute to downregulation of insulin signaling, resulting in loss of improved insulin sensitivity by postoperative week 9. Overall, our data identify a role for VDR in the preservation of insulin sensitivity after SG.

The fact that increased WAT and hepatic inflammation is present only in KO SG mice suggests that VDR plays an important immunoregulatory role following SG. However, the underlying mechanisms are unknown. KO SG inflammation develops despite surgically induced weight loss, which signifies that this VDR-dependent process is not induced by hypertrophied and hypoxic adipocytes as seen in obesity-associated inflammation (36). Vitamin D-VDR signaling blocks NF-κB and MAPK pathways, suppressing transcription of many pro-inflammatory genes in adipocytes and macrophages, including IL-6, IL-8, TNF-α and INF-γ (2, 7, 33). In this context, we hypothesize that SG in VDR KO mice may act as a ‘second hit’ to activate the pathological immune response with increased cytokine release and consequent monocyte recruitment to WAT. Increased infiltration of macrophages and other immune cells could potentiate pro-inflammatory cytokine release even further (7), ultimately leading to loss of insulin sensitivity in KO SG animals.

In addition to developing IR, KO SG mice demonstrated altered glucose utilization in response to oral challenge. It is possible that the delayed glucose clearance in KO SG animals represents impaired pancreatic insulin secretion secondary to lack of functional VDR (37) and low levels of calcium (38, 39). Additionally, without VDR, hepatic glucose production following an oral glucose load, might not be suppressed as efficiently as in WT animals due to hepatic insulin resistance (7, 40). Finally, worsened glucose regulation in KO SG mice could be related to low levels of glucoregulatory BA CA7S (15, 16). In our prior work we showed that with decreased levels of gastrointestinal LCA, the absorption of LCA into portal vein and its delivery to the liver are enhanced (15). Consequently, LCA induces CA7S production in the liver by binding to VDR and inducing SULT1A (15). Lack of VDR in KO SG mice compromised this pathway, providing a possible explanation for the decreased CA7S levels in these animals and their worsened glucose regulation as compared to WT SG animals.

However, the observation that CA7S levels were the highest in KO SH mice is surprising, as we have previously shown low to undetectable levels of CA7S in surgery-naïve VDR KO mice (15). It is possible that the surgical insult alone (without gastrectomy) induced changes resulting in higher CA7S levels in KO SH mice. Moreover, the modified total BA pool in KO SH mice could have blunted the antidiabetic effect of CA7S in these mice (15, 41). Further work will be needed to elucidate other pathways that might induce CA7S production and regulate its action in the host.

The only BA that was significantly higher in KO SG mice as compared to all the other groups was βMCA. βMCA is an antagonist of farnesoid X receptor (FXR) (42). Intestinal FXR antagonism has been linked with increased secretion of glucagon-like peptide 1 (GLP-1) which in turn potentiates insulin secretion (43). However, in KO SG mice we see the opposite phenotype, suggesting that the relatively high level of βMCA is offset by other factors, overall leading to worsened glucose regulation. On the other hand, βMCA is known to reduce intestinal absorption of cholesterol (44), suggesting a potential link between high levels of βMCA and low serum cholesterol in KO SG mice. Furthermore, one *in vitro* study suggested that βMCA can suppress lipid accumulation in primary mouse hepatocytes (45). Whether it reaches peripheral circulation and has the same effect in adipocytes, therefore contributing to their decreased size in KO SG mice, is unknown. Overall, further work is needed to determine the extent of βMCA contribution to the metabolic phenotype of KO SG mice.

We observed that over time, KO mice, regardless of surgery type, gain less weight than their WT counterparts. This is consistent with baseline characteristics of VDR KO strain: these mice fail to grow as rapidly as WT mice and exhibit approximately 10% lower body weight and decreased WAT mass – features that become more evident with ageing (19, 25). In this context, surgery in KO mice led to further decrease in adiposity, overall resulting in deficient WAT depots in KO SG mice as compared to WT SG mice. This observation, combined with loss of insulin sensitivity, suggests that KO SG mice exhibit lipodystrophy, defined as a deficiency or absence of AT, which is typically associated with IR (46). The underlying mechanism responsible for the exacerbated reduction in adiposity in KO SG animals is unknown, but it could involve an immune-mediated lysis of adipocytes as seen in certain forms of acquired lipodystrophies (46, 47). Alternatively, it could result from increased energy expenditure and fatty acid oxidation, which would also explain why there is no lipid accumulation in the liver or in the circulation in these mice (48). VDR KO mice are known to have increased energy expenditure (25, 28, 49), and it is possible that SG increases it further (50, 51), leading to exacerbated reduction in adiposity and hypolipidemia. Overall, our data demonstrate that without VDR, surgically induced reduction in AT mass is excessive and pathological, with resulting loss of normal tissue function.

Finally, we found that the expression of *VDR* gene is higher in SAT of individuals post MBS as compared to those without history of MBS. This provides evidence to support the claim that VDR is involved in the metabolic response to bariatric surgery in humans. Although studies on VDR expression post MBS are lacking, our data are in line with literature on vitamin D supplementation, which demonstrates positive effects of enhanced vitamin D – VDR signaling on metabolic health in humans. Increased VDR expression in response to vitamin D supplementation was shown to decrease body adiposity (11), improve hepatic insulin sensitivity and reduce AT inflammation (7). It is therefore possible that MBS induces expression of VDR in human AT, and via consequent enhanced vitamin D–VDR signaling, decreases AT inflammation and reduces insulin resistance.

Our study has some limitations. First, it is uncertain how the findings in an animal model translate to human physiology, especially in context of VDR-mediated regulation of body adiposity, where there is a known discrepancy between mouse and human data (28). Second, we only used male mice in our model, because they are more prone to DIO than females (52). Consequently, we cannot exclude the existence of potential sex differences in VDR-mediated responses to SG. Finally, further mechanistic studies are required to elucidate the exact pathways responsible for the unhealthy metabolic phenotype seen in KO SG mice.

In conclusion, we show that VDR is necessary for normal WAT function and glucose homeostasis after SG. Despite surgery-induced reduction in BW, KO SG mice demonstrated worsened metabolic health with altered glucose utilization, decrease in glucoregulatory CA7S, loss of insulin sensitivity, lipodystrophy, hypolipidemia, hypocalcemia, and immune dysregulation. Our study emphasizes the importance of VDR in maintaining metabolic benefits of MBS independent of weight loss.

## ACKNOWLEDGEMENT

Graphical abstract, Figure 1A and Figure 7A were created with BioRender.com (2025).

## FUNDING

This work was supported by National Institutes of Health (NIH) grant R01 DK126855 (E.G.S. and A.S.D.). Y.C. is supported by the American Heart Association Postdoctoral Fellowship (24POST1242610). C.F.R. received support from the NIH T32 Program for Research Training in Alimentary Tract Surgery (2T32DK007754–21).

## CONFLICT OF INTEREST

A.S.D. is an ad hoc consultant for Axial Therapeutics. E.G.S. has research support from Novo Nordisk and NIH, intellectual property related to diabetes and obesity treatment and has received past consulting/educational support from Vicarious Surgical, Intuitive, and Cine-Med. A.T. is a cofounder and consultant for AltrixBio.

## CRediT AUTHORSHIP CONTRIBUTION STATEMENT

**Andrei Moscalu**: Investigation, Data Curation, Formal Analysis; Writing – review & editing. **Weronika Stupalkowska**: Investigation, Data Curation, Formal Analysis, Visualization, Writing-original draft. **Lei Zhao**: Investigation, Formal Analysis, Writing-review & editing. **Yitong Li**: Investigation, Formal Analysis, Writing – review & editing. **Cullen F. Roberts**: Investigation, Data Curation, Writing-review & editing. **Yingjia Chen**: Investigation, Writing-review & editing. **Fei Ye**: Investigation, Formal Analysis, Writing-review & editing. **Rose Gold**: Data curation, Writing-review & editing. **Gavin Bewick**: Supervision, Writing-review & editing. **Francesco Rubino**: Supervision, Writing-review & editing. **A. Tavakkoli**: Supervision, Writing-review & editing. **A. Sloan Devlin**: Conceptualization, Supervision, Writing-review & editing, Funding acquisition. **Eric G. Sheu**: Conceptualization, Supervision, Writing-review & editing, Funding acquisition.

## DATA AVAILABILITY

Requests for further resources and data related to this paper should be directed to the corresponding authors (A.S.D. or E.G.S.).

## REFERENCES

1. Norman AW. Minireview: vitamin D receptor: new assignments for an already busy receptor. Endocrinology. 2006;147(12):5542–8.

2. Ding C, Gao D, Wilding J, Trayhurn P, Bing C. Vitamin D signalling in adipose tissue. Br J Nutr. 2012;108(11):1915–23.

3. Rosen CJ, Adams JS, Bikle DD, Black DM, Demay MB, Manson JE, et al. The nonskeletal effects of vitamin D: an Endocrine Society scientific statement. Endocr Rev. 2012;33(3):456–92.

4. Bassatne A, Chakhtoura M, Saad R, Fuleihan GE. Vitamin D supplementation in obesity and during weight loss: A review of randomized controlled trials. Metabolism. 2019;92:193–205.

5. Lei X, Zhou Q, Wang Y, Fu S, Li Z, Chen Q. Serum and supplemental vitamin D levels and insulin resistance in T2DM populations: a meta-analysis and systematic review. Sci Rep. 2023;13(1):12343.

6. Chen W, Liu L, Hu F. Efficacy of vitamin D supplementation on glycaemic control in type 2 diabetes: An updated systematic review and meta-analysis of randomized controlled trials. Diabetes Obes Metab. 2024;26(12):5713–26.

7. Lontchi-Yimagou E, Kang S, Goyal A, Zhang K, You JY, Carey M, et al. Insulin-sensitizing effects of vitamin D repletion mediated by adipocyte vitamin D receptor: Studies in humans and mice. Mol Metab. 2020;42:101095.

8. Argano C, Mirarchi L, Amodeo S, Orlando V, Torres A, Corrao S. The Role of Vitamin D and Its Molecular Bases in Insulin Resistance, Diabetes, Metabolic Syndrome, and Cardiovascular Disease: State of the Art. Int J Mol Sci. 2023;24(20).

9. Pieńkowska A, Janicka J, Duda M, Dzwonnik K, Lip K, Mędza A, et al. Controversial Impact of Vitamin D Supplementation on Reducing Insulin Resistance and Prevention of Type 2 Diabetes in Patients with Prediabetes: A Systematic Review. Nutrients. 2023;15(4).

10. Bouillon R, Manousaki D, Rosen C, Trajanoska K, Rivadeneira F, Richards JB. The health effects of vitamin D supplementation: evidence from human studies. Nat Rev Endocrinol. 2022;18(2):96–110.

11. Medeiros JFP, de Oliveira Borges MV, Soares AA, Dos Santos JC, de Oliveira ABB, da Costa CHB, et al. The impact of vitamin D supplementation on VDR gene expression and body composition in monozygotic twins: randomized controlled trial. Sci Rep. 2020;10(1):11943.

12. Xu Y, Lou Y, Kong J. VDR regulates energy metabolism by modulating remodeling in adipose tissue. Eur J Pharmacol. 2019;865:172761.

13. Kang S, Tsai LT, Zhou Y, Evertts A, Xu S, Griffin MJ, et al. Identification of nuclear hormone receptor pathways causing insulin resistance by transcriptional and epigenomic analysis. Nat Cell Biol. 2015;17(1):44–56.

14. Makishima M, Lu TT, Xie W, Whitfield GK, Domoto H, Evans RM, et al. Vitamin D receptor as an intestinal bile acid sensor. Science. 2002;296(5571):1313–6.

15. Chaudhari SN, Luo JN, Harris DA, Aliakbarian H, Yao L, Paik D, et al. A microbial metabolite remodels the gut-liver axis following bariatric surgery. Cell Host Microbe. 2021;29(3):408–24.e7.

16. Chaudhari SN, Harris DA, Aliakbarian H, Luo JN, Henke MT, Subramaniam R, et al. Bariatric surgery reveals a gut-restricted TGR5 agonist with anti-diabetic effects. Nat Chem Biol. 2021;17(1):20–9.

17. Eisenberg D, Shikora SA, Aarts E, Aminian A, Angrisani L, Cohen RV, et al. 2022 American Society of Metabolic and Bariatric Surgery (ASMBS) and International Federation for the Surgery of Obesity and Metabolic Disorders (IFSO) Indications for Metabolic and Bariatric Surgery. Obes Surg. 2023;33(1):3–14.

18. Zhang J, Feng M, Pan L, Wang F, Wu P, You Y, et al. Effects of vitamin D deficiency on the improvement of metabolic disorders in obese mice after vertical sleeve gastrectomy. Sci Rep. 2021;11(1):6036.

19. Li YC, Pirro AE, Amling M, Delling G, Baron R, Bronson R, et al. Targeted ablation of the vitamin D receptor: an animal model of vitamin D-dependent rickets type II with alopecia. Proc Natl Acad Sci U S A. 1997;94(18):9831–5.

20. Harris DA, Mina A, Cabarkapa D, Heshmati K, Subramaniam R, Banks AS, et al. Sleeve gastrectomy enhances glucose utilization and remodels adipose tissue independent of weight loss. Am J Physiol Endocrinol Metab. 2020;318(5):E678–e88.

21. Galarraga M, Campión J, Muñoz-Barrutia A, Boqué N, Moreno H, Martínez JA, et al. Adiposoft: automated software for the analysis of white adipose tissue cellularity in histological sections. J Lipid Res. 2012;53(12):2791–6.

22. Kleiner DE, Brunt EM, Van Natta M, Behling C, Contos MJ, Cummings OW, et al. Design and validation of a histological scoring system for nonalcoholic fatty liver disease. Hepatology. 2005;41(6):1313–21.

23. Yao L, Seaton SC, Ndousse-Fetter S, Adhikari AA, DiBenedetto N, Mina AI, et al. A selective gut bacterial bile salt hydrolase alters host metabolism. Elife. 2018;7.

24. Perez LJ, Rios L, Trivedi P, D’Souza K, Cowie A, Nzirorera C, et al. Validation of optimal reference genes for quantitative real time PCR in muscle and adipose tissue for obesity and diabetes research. Sci Rep. 2017;7(1):3612.

25. Narvaez CJ, Matthews D, Broun E, Chan M, Welsh J. Lean phenotype and resistance to diet-induced obesity in vitamin D receptor knockout mice correlates with induction of uncoupling protein-1 in white adipose tissue. Endocrinology. 2009;150(2):651–61.

26. Jahn D, Dorbath D, Schilling AK, Gildein L, Meier C, Vuille-Dit-Bille RN, et al. Intestinal vitamin D receptor modulates lipid metabolism, adipose tissue inflammation and liver steatosis in obese mice. Biochim Biophys Acta Mol Basis Dis. 2019;1865(6):1567–78.

27. Li YC, Amling M, Pirro AE, Priemel M, Meuse J, Baron R, et al. Normalization of mineral ion homeostasis by dietary means prevents hyperparathyroidism, rickets, and osteomalacia, but not alopecia in vitamin D receptor-ablated mice. Endocrinology. 1998;139(10):4391–6.

28. Bouillon R, Carmeliet G, Lieben L, Watanabe M, Perino A, Auwerx J, et al. Vitamin D and energy homeostasis: of mice and men. Nat Rev Endocrinol. 2014;10(2):79–87.

29. Rafiq S, Jeppesen PB. Vitamin D Deficiency Is Inversely Associated with Homeostatic Model Assessment of Insulin Resistance. Nutrients. 2021;13(12).

30. Han FF, Lv YL, Gong LL, Liu H, Wan ZR, Liu LH. VDR Gene variation and insulin resistance related diseases. Lipids Health Dis. 2017;16(1):157.

31. Maestro B, Molero S, Bajo S, Dávila N, Calle C. Transcriptional activation of the human insulin receptor gene by 1,25-dihydroxyvitamin D(3). Cell Biochem Funct. 2002;20(3):227–32.

32. Manna P, Achari AE, Jain SK. 1,25(OH)(2)-vitamin D(3) upregulates glucose uptake mediated by SIRT1/IRS1/GLUT4 signaling cascade in C2C12 myotubes. Mol Cell Biochem. 2018;444(1-2):103–8.

33. Nimitphong H, Guo W, Holick MF, Fried SK, Lee MJ. Vitamin D Inhibits Adipokine Production and Inflammatory Signaling Through the Vitamin D Receptor in Human Adipocytes. Obesity (Silver Spring). 2021;29(3):562–8.

34. Marcotorchino J, Gouranton E, Romier B, Tourniaire F, Astier J, Malezet C, et al. Vitamin D reduces the inflammatory response and restores glucose uptake in adipocytes. Mol Nutr Food Res. 2012;56(12):1771–82.

35. Schmid A, Karrasch T, Schäffler A. The emerging role of bile acids in white adipose tissue. Trends Endocrinol Metab. 2023;34(11):718–34.

36. Saltiel AR, Olefsky JM. Inflammatory mechanisms linking obesity and metabolic disease. J Clin Invest. 2017;127(1):1–4.

37. Wang X, Zhao X, Zhou R, Gu Y, Zhu X, Tang Z, et al. Delay in glucose peak time during the oral glucose tolerance test as an indicator of insulin resistance and insulin secretion in type 2 diabetes patients. J Diabetes Investig. 2018;9(6):1288–95.

38. Yasuda K, Hurukawa Y, Okuyama M, Kikuchi M, Yoshinaga K. Glucose tolerance and insulin secretion in patients with parathyroid disorders. Effect of serum calcium on insulin release. N Engl J Med. 1975;292(10):501–4.

39. Chang E, Donkin SS, Teegarden D. Parathyroid hormone suppresses insulin signaling in adipocytes. Mol Cell Endocrinol. 2009;307(1-2):77–82.

40. Oh J, Riek AE, Darwech I, Funai K, Shao J, Chin K, et al. Deletion of macrophage Vitamin D receptor promotes insulin resistance and monocyte cholesterol transport to accelerate atherosclerosis in mice. Cell Rep. 2015;10(11):1872–86.

41. Chen Y, Chaudhari SN, Harris DA, Roberts CF, Moscalu A, Mathur V, et al. A small intestinal bile acid modulates the gut microbiome to improve host metabolic phenotypes following bariatric surgery. Cell Host Microbe. 2024;32(8):1315–30.e5.

42. Sayin SI, Wahlström A, Felin J, Jäntti S, Marschall HU, Bamberg K, et al. Gut microbiota regulates bile acid metabolism by reducing the levels of tauro-beta-muricholic acid, a naturally occurring FXR antagonist. Cell Metab. 2013;17(2):225–35.

43. Trabelsi MS, Daoudi M, Prawitt J, Ducastel S, Touche V, Sayin SI, et al. Farnesoid X receptor inhibits glucagon-like peptide-1 production by enteroendocrine L cells. Nat Commun. 2015;6:7629.

44. Wang DQ, Tazuma S. Effect of beta-muricholic acid on the prevention and dissolution of cholesterol gallstones in C57L/J mice. J Lipid Res. 2002;43(11):1960–8.

45. Takada S, Matsubara T, Fujii H, Sato-Matsubara M, Daikoku A, Odagiri N, et al. Stress can attenuate hepatic lipid accumulation via elevation of hepatic β-muricholic acid levels in mice with nonalcoholic steatohepatitis. Lab Invest. 2021;101(2):193–203.

46. Brown RJ, Araujo-Vilar D, Cheung PT, Dunger D, Garg A, Jack M, et al. The Diagnosis and Management of Lipodystrophy Syndromes: A Multi-Society Practice Guideline. J Clin Endocrinol Metab. 2016;101(12):4500–11.

47. Mandel-Brehm C, Vazquez SE, Liverman C, Cheng M, Quandt Z, Kung AF, et al. Autoantibodies to Perilipin-1 Define a Subset of Acquired Generalized Lipodystrophy. Diabetes. 2023;72(1):59–70.

48. Reue K, Phan J. Metabolic consequences of lipodystrophy in mouse models. Curr Opin Clin Nutr Metab Care. 2006;9(4):436–41.

49. Wong KE, Szeto FL, Zhang W, Ye H, Kong J, Zhang Z, et al. Involvement of the vitamin D receptor in energy metabolism: regulation of uncoupling proteins. Am J Physiol Endocrinol Metab. 2009;296(4):E820–8.

50. McGavigan AK, Garibay D, Henseler ZM, Chen J, Bettaieb A, Haj FG, et al. TGR5 contributes to glucoregulatory improvements after vertical sleeve gastrectomy in mice. Gut. 2017;66(2):226–34.

51. Li P, Rao Z, Laing BT, Bunner W, Landry T, Prete A, et al. Vertical sleeve gastrectomy improves liver and hypothalamic functions in obese mice. J Endocrinol. 2019;241(2):135–47.

52. Casimiro I, Stull ND, Tersey SA, Mirmira RG. Phenotypic sexual dimorphism in response to dietary fat manipulation in C57BL/6J mice. J Diabetes Complications. 2021;35(2):107795.

