## Supplemental File for "Vitamin D receptor is necessary for metabolic health after sleeve gastrectomy"

**Table S1.** Composition of rodent diet used in this study.

| Ingredient | Gram | Kcal |
|---|---|---|
| Casein | 200 | 800 |
| L-Cystine | 3 | 12 |
| Maltodextrin 10 | 32.71 | 131 |
| Sucrose | 0 | 0 |
| Lactose | 161.09 | 644 |
| Cellulose | 50 | 0 |
| Soybean Oil | 25 | 225 |
| Lard | 245 | 2205 |
| Mineral Mix, S10026 | 10 | 0 |
| DiCalcium Phosphate | 38 | 0 |
| Calcium Carbonate | 12.1 | 0 |
| Potassium Citrate | 16.5 | 0 |
| Vitamin Mix V10001 | 10 | 40 |
| Choline Bitartrate | 2 | 0 |
| FD&C Blue Dye | 0.05 | 0 |
| **Total** | **805.45** | **4057** |

**Table S2.** Scoring system used in this study to assess adipose tissue inflammation, necrosis and fibrosis.

| Score | Description | |
|---|---|---|
| | ***Inflammation*** | ***Necrosis*** |
| 0 | No inflammatory cells | No necrosis |
| 1 | Minimal inflammatory infiltrate | Few minimal foci of necrosis |
| 2 | Moderate inflammatory infiltrate | Several foci of necrosis |
| 3 | Severe inflammatory infiltrate | Extensive necrosis |

**Table S3.** Primer sequences used for qPCR in this study.

| Gene | Primer | Identifier |
|---|---|---|
| Human *VDR* | F - GCCGGACCAGAAGCCTTT | This study |
| | R - TCTCCACACACCCCACAGAT | |
| Human *HSPCB* | F – TTATTTTAGATGCCTGAGGAAGTGC | This study |
| | R- CGAATCTTGTCCAAGGCATCAG | |

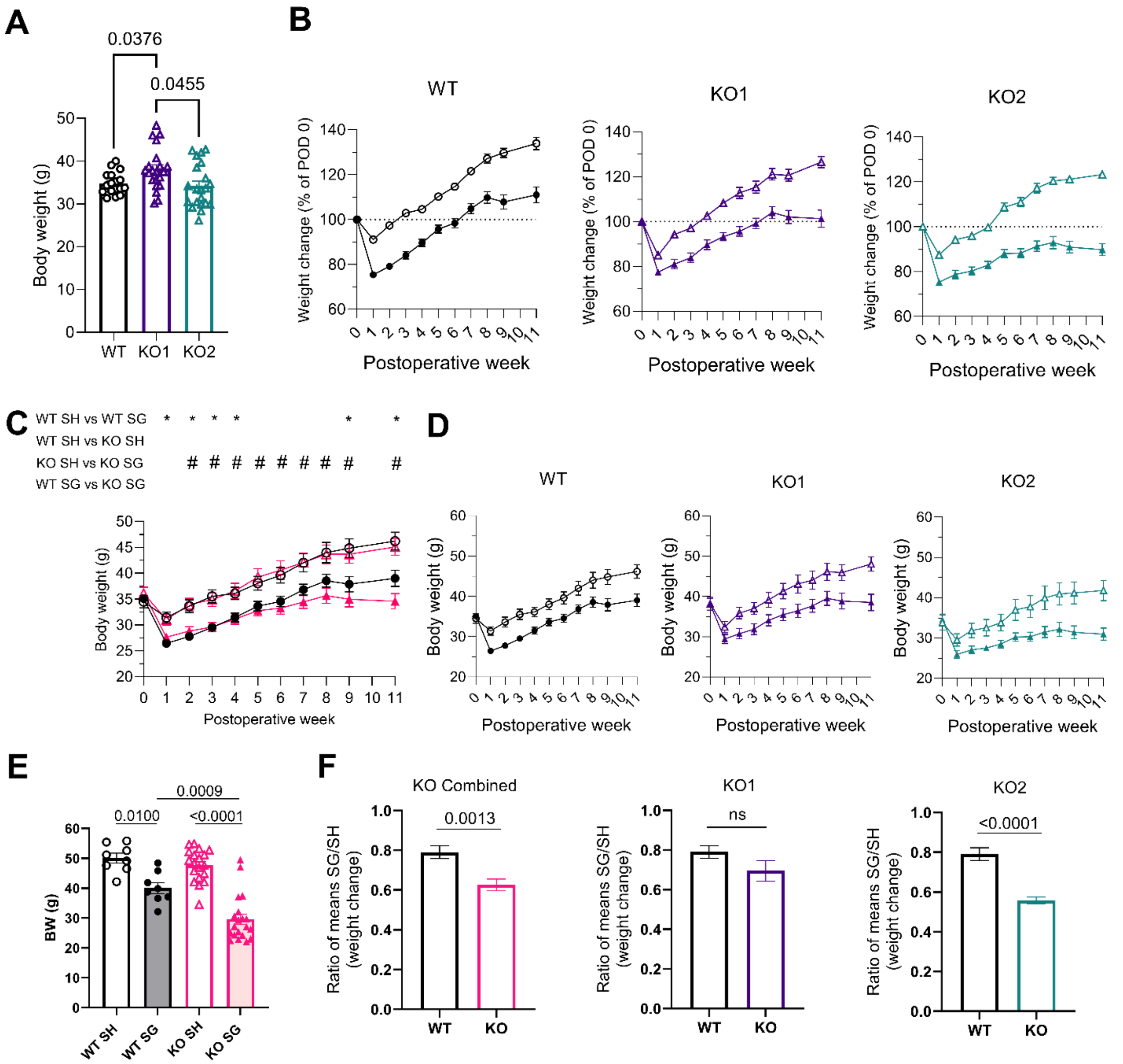


**Figure S1.** (A) Preoperative body weight for WT cohort and two VDR knockout cohorts. (B) Change in body weight over time as % of body weight on postoperative day 0 (POD 0) plotted for WT cohort and KO cohorts separately. (C) Absolute body weight over time. (D) Absolute body weight for WT cohort and KO cohort separately. (E) Absolute body weights on necropsy day. (F) SG/SH ratio of mean weight change on necropsy day plotted for KO cohorts combined (left) and separately (middle and right). Animal numbers: WT SH (n=8), WT SG (n=9), KO1 SH (n=11), KO1 SG (n=10), KO2 SH (n=10), KO2 SG (n=11). Data in panel A were analyzed with one way ANOVA with Dunnett's multiple comparisons test. Data in panel C were analyzed with repeated measures two-way ANOVA with Tukey's multiple comparisons test and corresponding p values are given in Table S3. Data in panel E were analyzed with two-way ANOVA with Sidak's multiple comparisons test. Data in panel F were analyzed with Student's t-tests.

**Table S2**. P values for Tukey's multiple comparisons test after two-way repeated measures ANOVA in Figure 1, panel B.

| Week | Comparison | | | |
|---|---|---|---|---|
| | WT SH vs WT SG * | WT SH vs KO SH ^ | KO SH vs KO SG # | WT SG vs KO SG $ |
| 1 | <0.0001 | 0.0063 | <0.0001 | ns |
| 2 | <0.0001 | 0.0444 | <0.0001 | ns |
| 3 | <0.0001 | <0.0001 | <0.0001 | ns |
| 4 | <0.0001 | 0.0207 | <0.0001 | ns |
| 5 | 0.0001 | ns | <0.0001 | ns |
| 6 | 0.0001 | ns | <0.0001 | ns |
| 7 | 0.0001 | ns | <0.0001 | 0.0136 |
| 8 | 0.0005 | ns | <0.0001 | 0.0134 |
| 9 | 0.0003 | 0.0169 | <0.0001 | 0.0422 |
| 11 | 0.0006 | ns | <0.0001 | 0.0102 |

**Table S3**. P values for Tukey's multiple comparisons test after two-way repeated measures ANOVA in Figure S2, panel C.

| Week | Comparison | | | |
|---|---|---|---|---|
| | WT SH vs WT SG * | WT SH vs KO SH ^ | KO SH vs KO SG # | WT SG vs KO SG $ |
| 1 | 0.0070 | ns | ns | ns |
| 2 | 0.0073 | ns | 0.0066 | ns |
| 3 | 0.0082 | ns | 0.0102 | ns |
| 4 | 0.0363 | ns | 0.0212 | ns |
| 5 | ns | ns | 0.0097 | ns |
| 6 | ns | ns | 0.0090 | ns |
| 7 | ns | ns | 0.0074 | ns |
| 8 | ns | ns | 0.0071 | ns |
| 9 | 0.0462 | ns | 0.0023 | ns |
| 11 | 0.0343 | ns | 0.0001 | ns |

**Table S4**. P values for Tukey's multiple comparisons test after two-way repeated measures ANOVA in Figure 1, panel D.

| Week | Comparison | | | |
|---|---|---|---|---|
| | WT SH vs WT SG * | WT SH vs KO SH ^ | KO SH vs KO SG # | WT SG vs KO SG $ |
| 1 | 0.0121 | 0.0171 | <0.0001 | ns |
| 2 | ns | 0.0153 | ns | Ns |
| 3 | ns | 0.0382 | ns | 0.0069 |
| 4 | ns | ns | ns | 0.0473 |
| 5 | ns | ns | ns | ns |
| 6 | ns | 0.0086 | ns | 0.0010 |
| 7 | ns | 0.0462 | ns | 0.0043 |
| 8 | ns | 0.0049 | ns | ns |
| 9 | ns | ns | ns | 0.0386 |

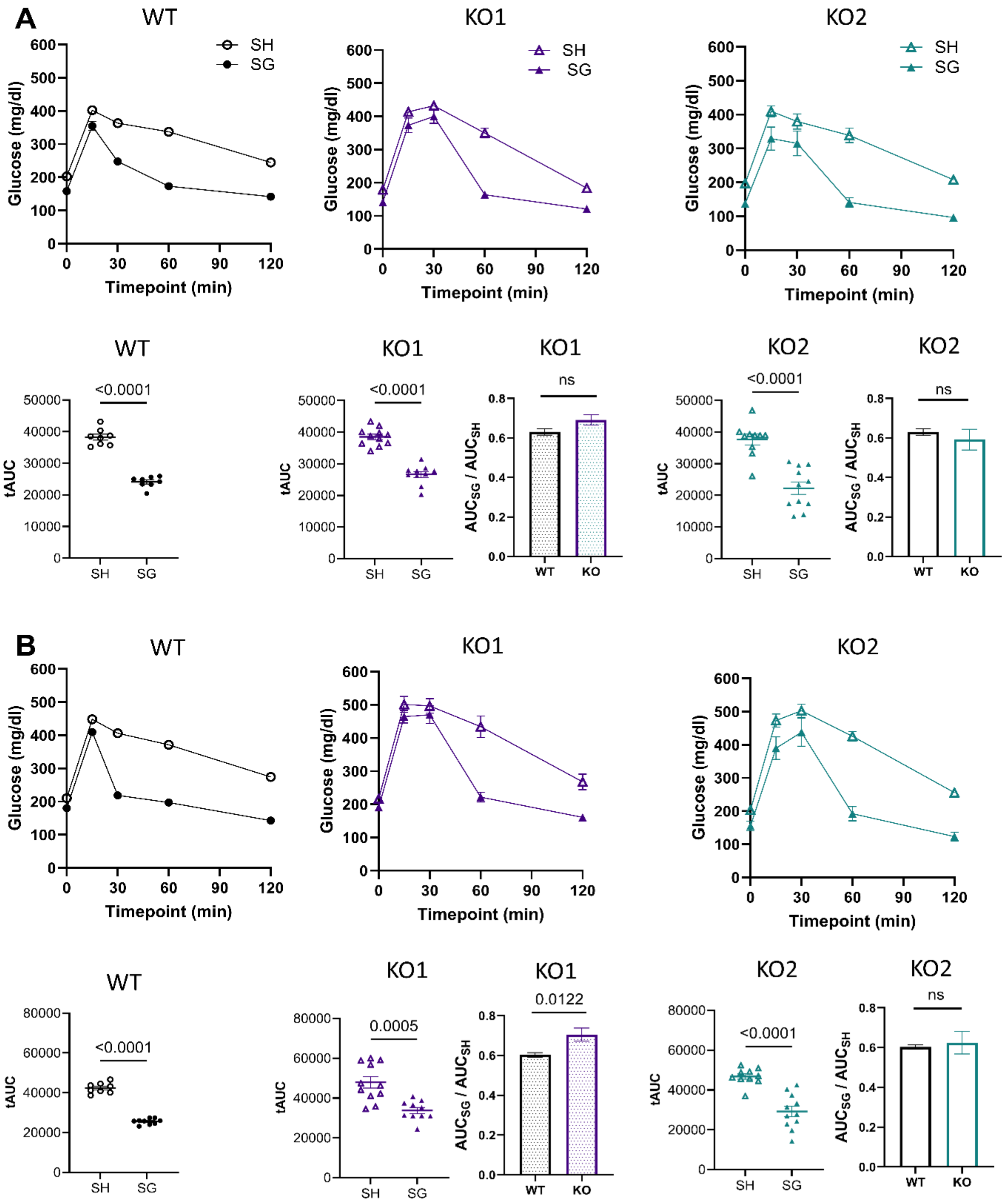


**Figure S2**. 2-week (A) and 8-week (B) oral glucose tolerance test with total AUC per experimental group and SG/SH ratios of mean AUC values plotted separately for WT cohort and KO cohorts. Animal numbers: WT SH (n=8), WT SG (n=9), KO1 SH (n=11), KO1 SG (n=10), KO2 SH (n=10), KO2 SG (n=11). In panel A one WT SG mouse was excluded from the analysis due to failure of oral gavage. Data are shown as mean ± SEM. AUC data were analyzed with two-tailed Student's t tests.

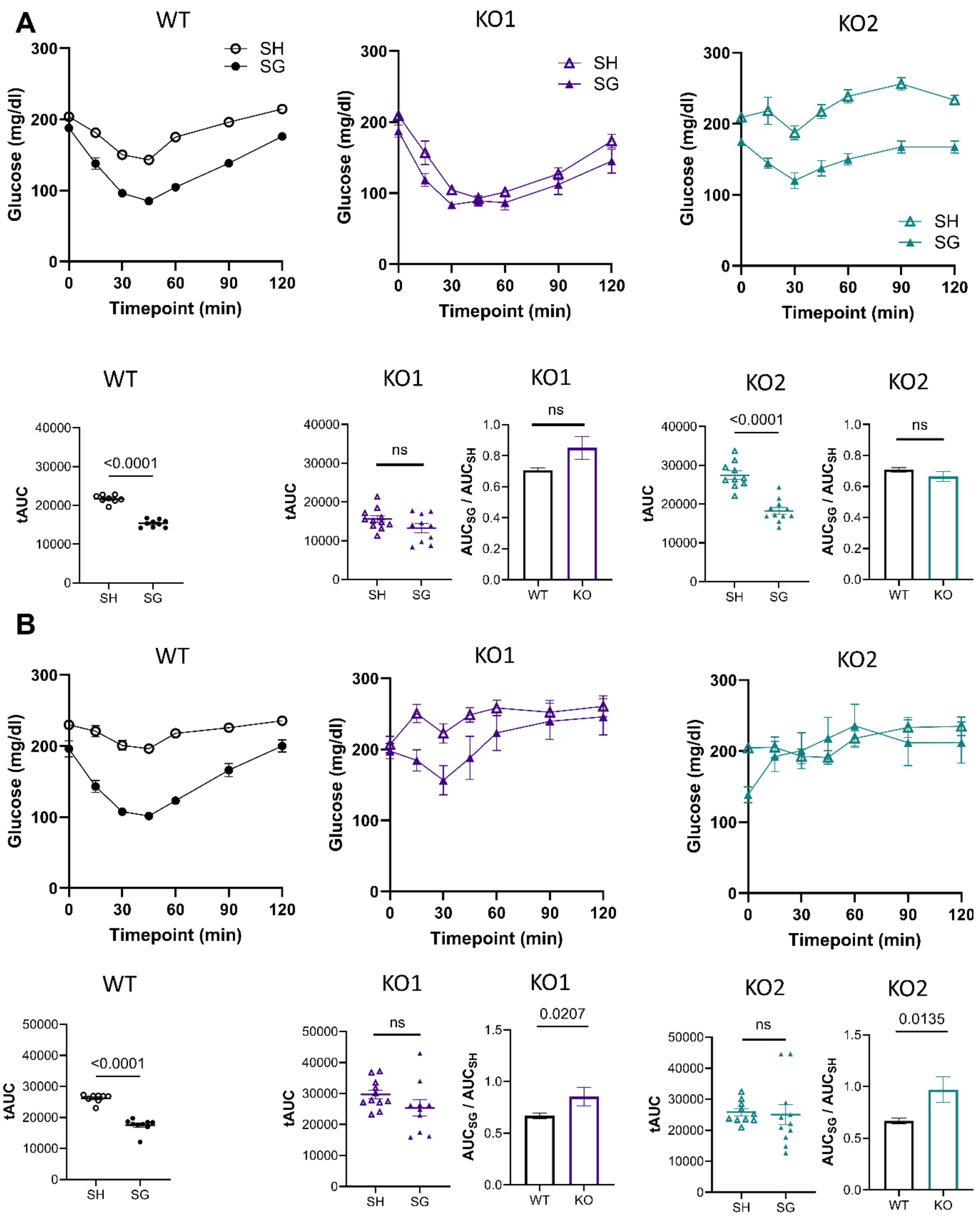


**Figure S3**. 5-week (A) and 9-week (B) insulin tolerance test with total AUC per experimental group and SG/SH ratios of mean AUC values plotted separately for WT cohort and KO cohorts. Animal numbers: WT SH (n=8), WT SG (n=9), KO1 SH (n=11), KO1 SG (n=10), KO2 SH (n=10), KO2 SG (n=11). Data are shown as mean ± SEM. AUC data were analyzed with two-tailed Student's t tests.

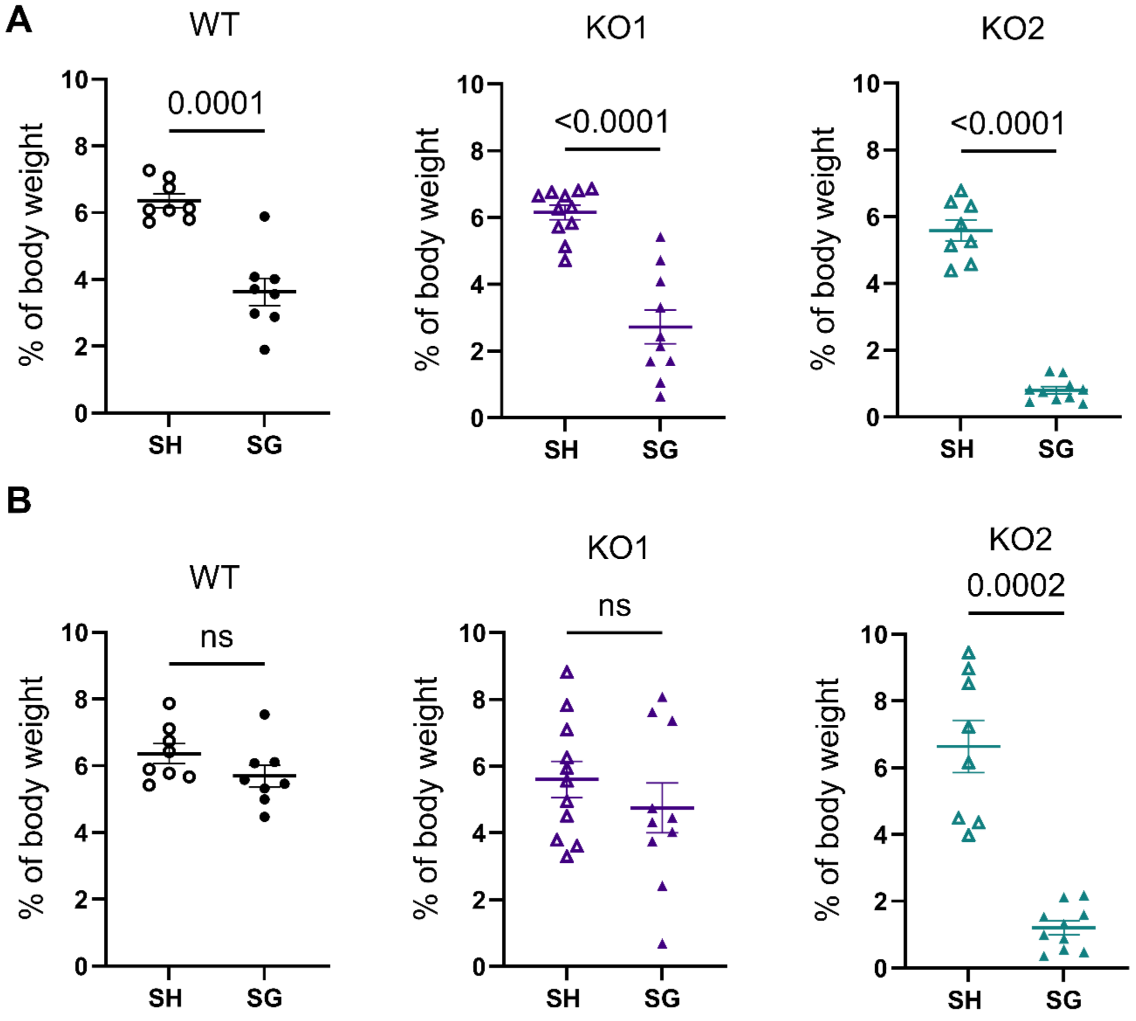


**Figure S4.** (A) Mass of inguinal (iWAT) and (B) epididymal (eWAT) white adipose tissue depots as percentage of body weight on necropsy day plotted separately for WT and KO cohorts. Animal numbers: WT SH (n=8), WT SG (n=8), KO1 SH (n=11), KO1 SG (n=10), KO2 SH (n=8), KO2 SG (n=10). Data are shown as mean ± SEM. Data were analyzed with two-tailed Student’s t test.

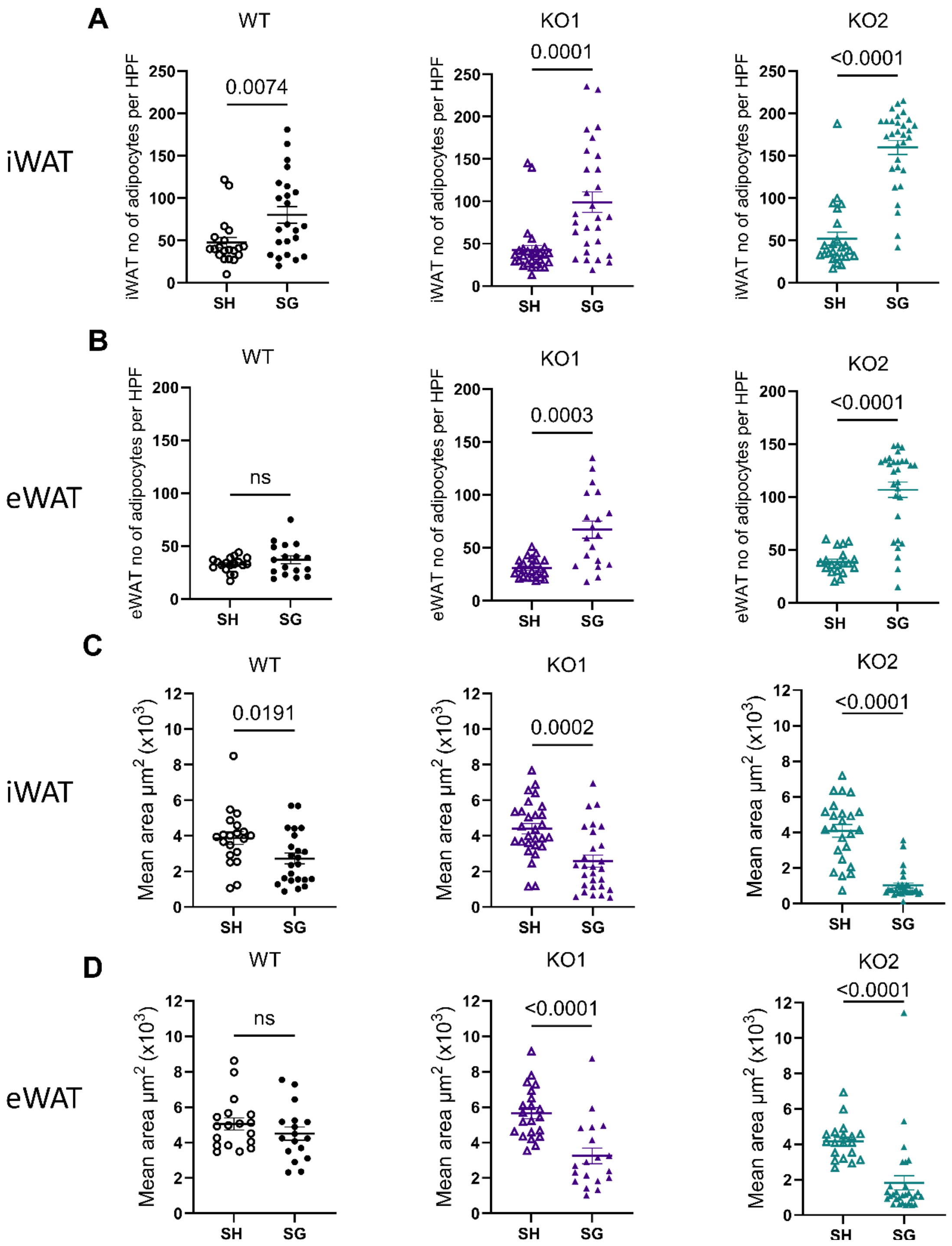


**Figure S5.** Number of adipocytes per x10 high power field (HPF) in iWAT (A) and eWAT (B) plotted separately for WT and KO cohorts. Mean adipocyte area in iWAT (C) and eWAT (D) plotted separately for WT and KO cohorts. Animal numbers: WT SH (n=8), WT SG (n=8), KO1 SH (n=11), KO1 SG (n=10), KO2 SH (n=8), KO2 SG (n=10). Each individual symbol represents an individual photomicrograph (for each mouse, one H&E slide was produced per adipose tissue type and 1-3 representative photomicrographs were taken of each slide for automated analysis). Data are shown as mean ± SEM. Data were analyzed with two-tailed Student's t test.

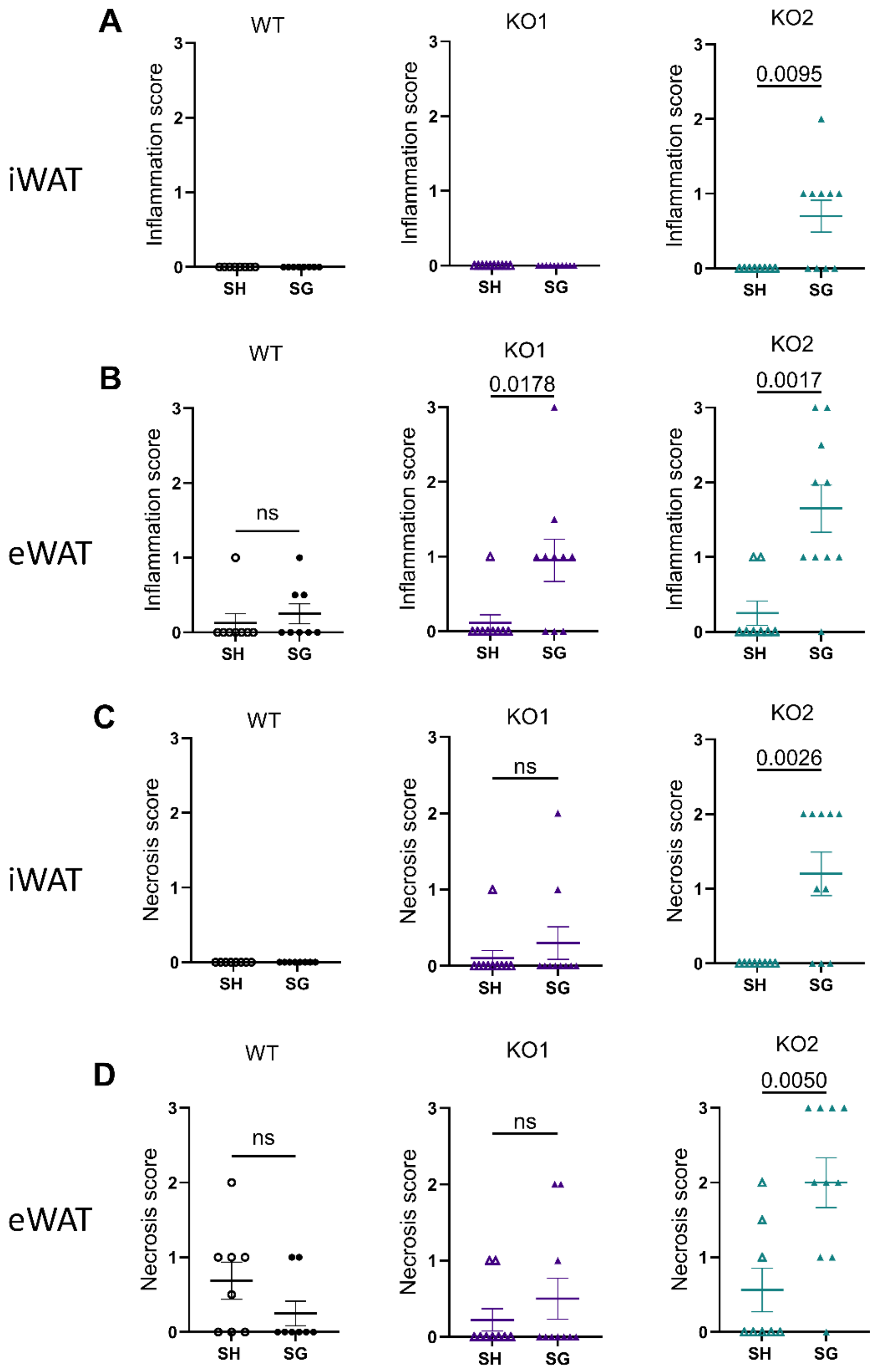


**Figure S6.** Inflammation score (0-3) in eWAT (A) and iWAT (B) plotted separately for WT and KO cohorts. Fat necrosis score (0-3) in eWAT (C) and iWAT (D) plotted separately for WT and KO cohorts. Animal numbers: WT SH (n=8), WT SG (n=8), KO1 SH (n=11), KO1 SG (n=10), KO2 SH (n=8), KO2 SG (n=10). Data are shown as mean ± SEM. Data were analyzed with two-tailed Student's t test. Data not marked indicate that t-test was not computed since all values are identical.

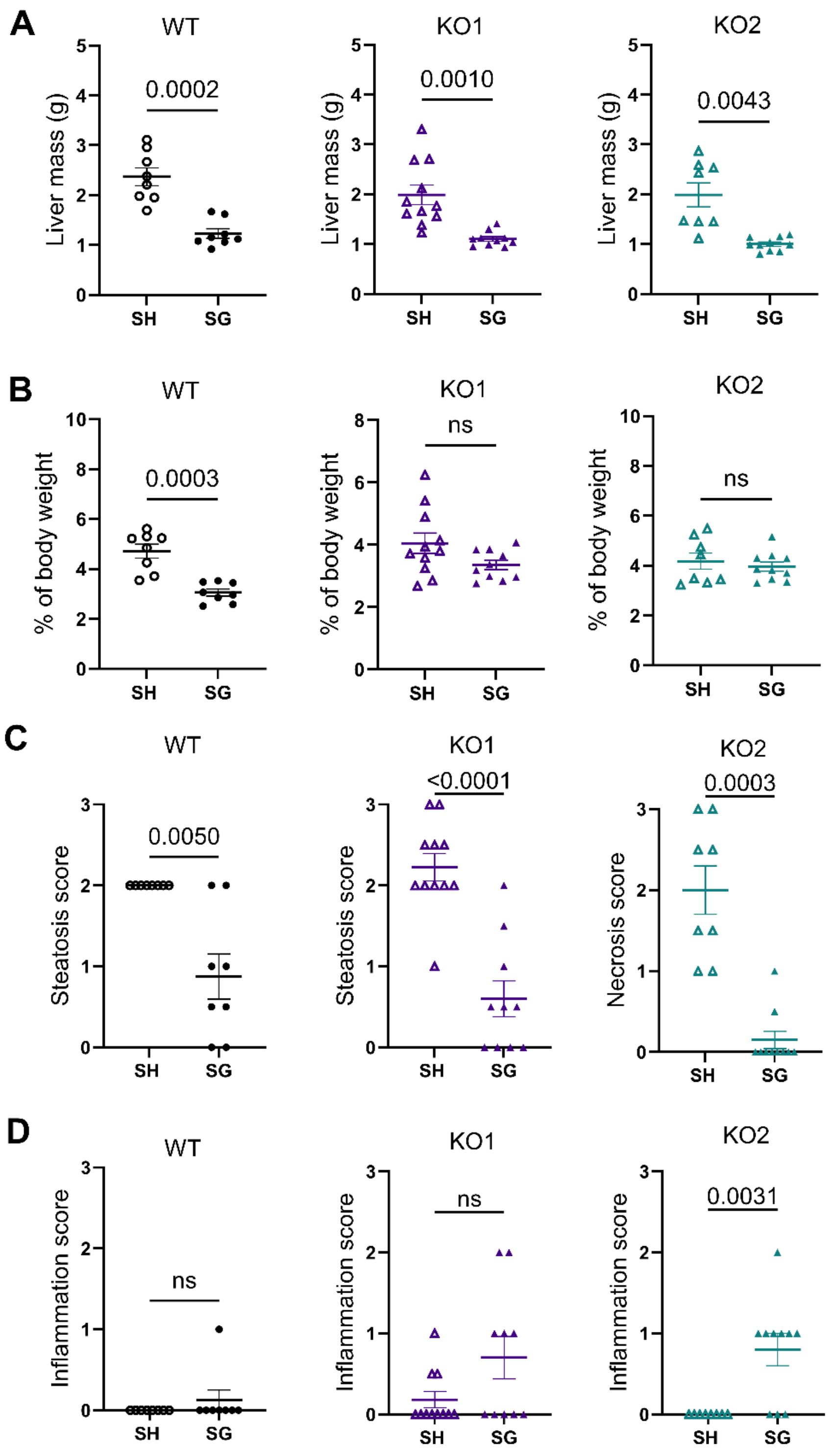


**Figure S7.** Absolute liver mass (A) and liver mass as percentage of body weight (B) on necropsy day plotted separately for WT and KO cohorts. Hepatic steatosis (C) and inflammation (D) scores (0-none, 1-mild, 2-moderate, 3-severe) plotted separately for WT and KO cohorts. Animal numbers: WT SH (n=8), WT SG (n=8), KO1 SH (n=11), KO1 SG (n=10), KO2 SH (n=8), KO2 SG (n=10). Data are shown as mean ± SEM. Data were analyzed with two-tailed Student's t test.

**Table S5.** Cases of liver fibrosis in all experimental groups as assessed on histology (H&E and Sirius Red stain).

| Experimental Group | LIVER FIBROSIS | |
|---|---|---|
| | PRESENT | ABSENT |
| WT SH | 0 | 8 |
| WT SG | 0 | 8 |
| KO SH | 0 | 19 |
| KO SG | 4 | 16 |
| Fisher's exact test | P value = 0.1230 | |

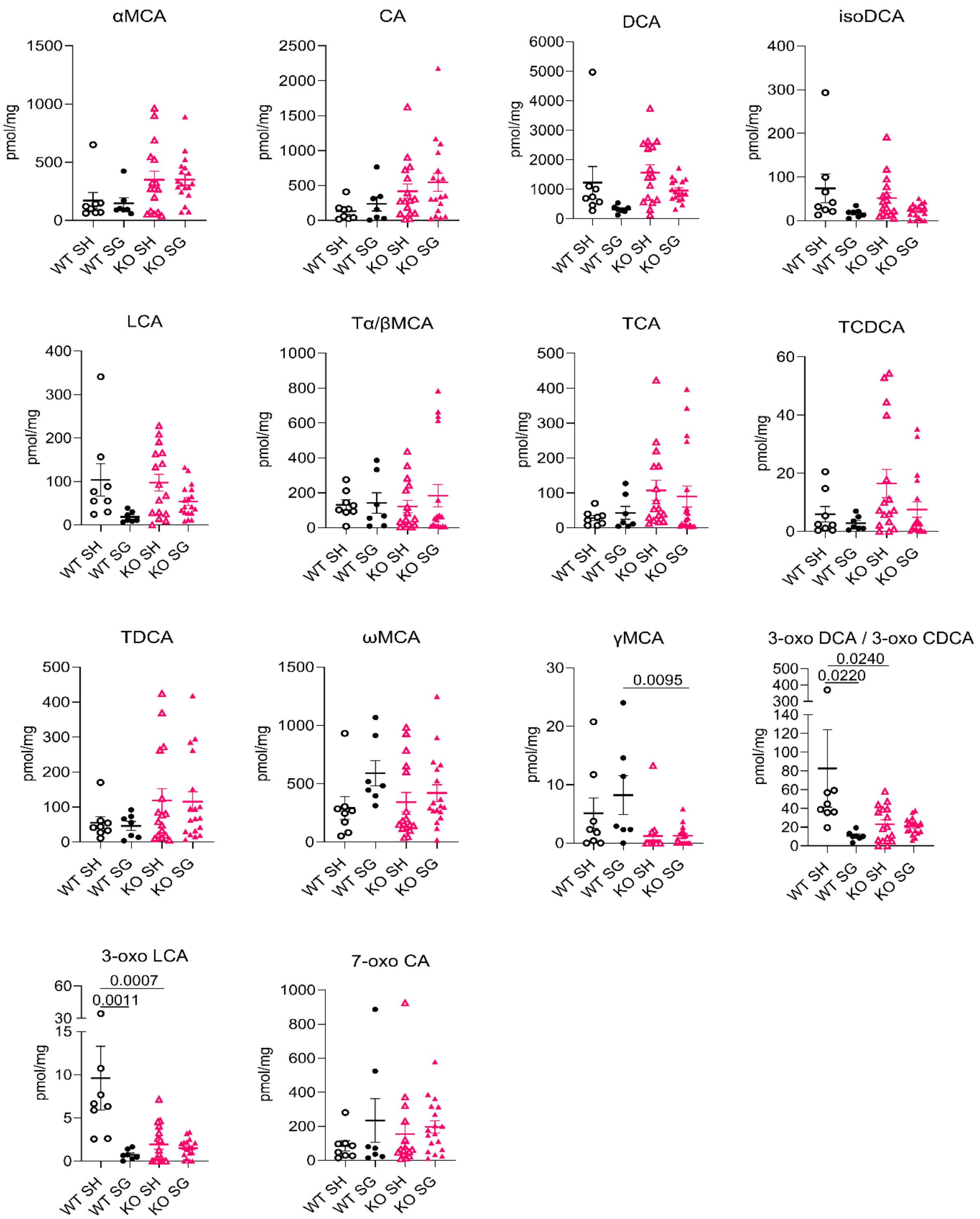


**Figure S8**. Individual bile acid concentrations in colonic contents of WT and VDR KO mice post-sham and post SG. All bile acids except CA7S and βMCA (depicted in Fig 6) with measurable concentrations above the limit of detection are included. Animal numbers: WT SH (n=8), WT SG (n=7), KO SH (n=16), KO SG (n=18). Each symbol represents an individual mouse. Data

presented as mean ± SEM. Data was analyzed with two-way ANOVA with Sidak's multiple comparisons test. Data not marked are not significant.
*Abbreviations: αMCA, alpha-muricholic acid; CA, cholic acid; DCA, deoxycholic acid; isoDCA, isodeoxycholic acid; LCA, lithocholic acid; Tα/βMCA, tauro-alpha and tauro-beta-muricholic acid*; *TCA, taurocholic acid; TCDCA, taurochenodeoxycholic acid; TDCA, taurodeoxycholic acid; ωMCA, omega-muricholic acid; 3-oxo DCA / 3-oxo CDCA; 3-oxo deoxycholic acid and 3-oxo chenodeoxycholic acid; 3-oxo LCA, 3-oxo lithocholic acid; 7-oxo CA, 7-oxo cholic acid; CA7S, cholic acid 7-sulphate; βMCA, beta-muricholic acid.*
